# Under-Oil Autonomously Regulated Oxygen Microenvironments: A Goldilocks Principle-Based Approach For Microscale Cell Culture

**DOI:** 10.1101/2020.12.16.423117

**Authors:** Chao Li, Mouhita Humayun, Glenn M. Walker, Keon Young Park, Bryce Connors, Jun Feng, Molly C. Pellitteri Hahn, Cameron O. Scarlett, Jiayi Li, Yanbo Feng, Ryan L. Clark, Hunter Hefti, Jonathan Schrope, Ophelia S. Venturelli, David J. Beebe

## Abstract

Oxygen levels *in vivo* are autonomously regulated by a supply-demand balance, which can be altered in disease states. However, the oxygen levels of *in vitro* cell culture systems, particularly microscale cell culture, are typically dominated by either supply or demand. Further, the oxygen microenvironment in these systems are rarely monitored or reported. Here, we present a method to establish and dynamically monitor autonomously regulated oxygen microenvironments (AROM) using an oil overlay in an open microscale cell culture system. Using this method, the oxygen microenvironment is dynamically regulated via a supply-demand balance of the system. We simulate the kinetics of oxygen diffusion in multiliquid-phase microsystems on COMSOL Multiphysics and experimentally validate the method using a variety of cell types including mammalian, fungal and bacterial cells. Finally, we demonstrate the utility of this method to establish a co-culture between primary intestinal epithelial cells and a highly prevalent human gut species *Bacteroides uniformis*.

## 1. Introduction

In the body, cells consume oxygen diffusing from capillaries nearby and continuously regulate and respond to their oxygen microenvironment.^[1]^ The oxygen microenvironment influences cellular and tissue functions of normal and disease states, where the local oxygen levels are defined by the pericellular oxygen concentration (POC) and intracellular oxygen concentration (IOC) (**Figure 1**a). *In vivo*, oxygen levels regulate diverse cellular activities and disease states including stem-cell states,^[2]^ stem-cell differentiation,^[3]^ immune response in inflammation,^[4,5]^ and pathogenesis of diseases such as cancer^[6]^ and inflammatory bowel disease.^[7]^ The oxygen microenvironment in local tissue is continuously and dynamically regulated by a supply-demand balance. Homeostasis is achieved when oxygen (O_2_) flux from the supply (i.e. oxygen carried in capillaries) matches O_2_ demand of the cells through the diffusion barrier (i.e. extracellular matrix, or ECM), and changes in these parameters alter the oxygen microenvironment. *In vivo*, changes in O_2_ supply, for instance, can be caused by fluctuating blood flow in the chaotic and poorly hierarchical structure of tumor blood vessels.^[8]^ Similarly, O_2_ demands can change during activation/repression of inflammation^[9]^ or during a switch between bioenergetic profiles.^[10]^ For example, a hallmark of inflammation and many disease states is hypoxia caused by impaired supply and/or upregulated demand of oxygen relative to physiologically defined oxygen levels, also known as physioxia. For a given supply and demand of oxygen, a Goldilocks or “just right” diffusion barrier is necessary to reach physioxia in the system (Figure 1b). *In vivo*, hypoxia or anoxia can be also caused by decreased tissue perfusion due to fibrosis.^[11]^ These physiological/pathological changes cause variations in POC and IOC leading to aberrant oxygen levels that further elicit changes in cell function and cell-to-cell interactions.^[12]^

**Figure 1.**
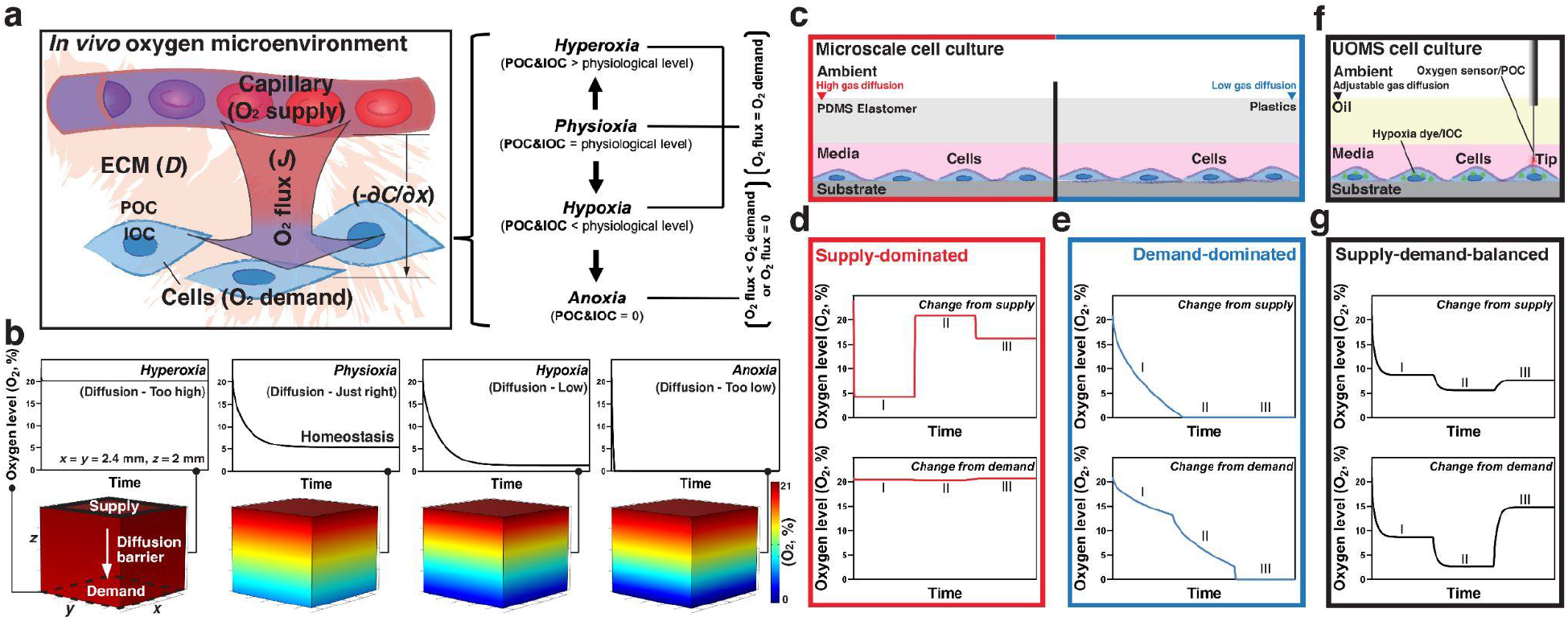
Oxygen microenvironments in *in vivo* and *in vitro* microscale cell culture. a) Schematic of a typical *in vivo* cellular and oxygen microenvironment. Different oxygen levels including hyperoxia, physioxia, hypoxia, and anoxia are defined by POC and IOC compared to the normal or physiological level of oxygen concentration *in vivo* (about 5% O_2_ in average). Hyperoxia, physioxia, or hypoxia is a result of homeostasis between O_2_ flux and O_2_ demand (i.e. O_2_ flux = O_2_ demand). In case of O_2_ flux is less than O_2_ demand homeostasis cannot be reached and the system ends up with anoxia. b) COMSOL Multiphysics simulation (Experimental Section) shows the typical kinetics of oxygen diffusion against varying diffusion barriers. Oxygen (21% O_2_) was set to diffuse from the supply (the top surface denoted by the black solid-line box on the model) to the demand (the bottom surface denoted by the black dashed-line box on the model) in the geometry of a standard 384 well (i.e. *x* = *y* = 2.4 mm). The diffusion barrier was set 2 mm in depth (i.e. *z* = 2 mm). The walls and the bottom surface (or the substrate) were defined as gas impermeable. The readout of oxygen level (O_2_, %) was extracted from the cell layer at the bottom surface. c) Schematic of typical (closed-channel or closed-chamber) microscale cell culture systems. The oxygen microenvironment in these microscale devices is either supply-dominated (left), due to the high gas diffusivity of PDMS elastomer and small scale, or demand-dominated (right) when materials (e.g. PS or PMMA) with limited gas diffusivity are used. d) and e) COMSOL Multiphysics results show the typical kinetics of oxygen level (at the cell layer) against the change from supply or demand with three stages (I, II, and III) in supply-dominated systems (the red box) and demand-dominated systems (the blue box). A supply-dominated system is highly responsive to the change from supply (i.e. external settings of oxygen) but leaves the demand (i.e. oxygen consumption of cells) largely disconnected from regulating the oxygen microenvironment. By contrast, a demand-dominated system responds little to the change from supply and leads to anoxia (i.e. complete depletion of oxygen at the cell layer) and eventually cell death. f) Schematic shows UOMS cell culture. Cells are cultured in media with an oil overlay - a proper diffusion barrier that allows a mimicry of the supply-demand-balanced oxygen microenvironment *in vivo*. For a given ambient and cell-media condition, gas diffusion through the oil overlay can be readily adjusted by selecting different oil properties (e.g. oil type, depth, viscosity). Enabled by the free physical access of UOMS, POC can be monitored with high spatial flexibility using an optical oxygen sensor deployed in contact to the cell layers through the oil overlay. IOC is probed by a hypoxia dye (depicted as green triangles) in live cells. g) COMSOL Multiphysics results show the typical kinetics of oxygen level (at the cell layer) to the change from supply or demand in a supply-demand-balanced system (the black box). A supply-demand-balanced system is able to respond to changes from both supply and demand and reaches to different stages of homeostasis autonomously.

Compared to conventional, bulk-scale cell culture techniques (e.g. culture flasks), microscale cell culture (i.e. low volume ratio between media and cells with the media volume typically falling in the range of picoliters to microliters and commonly < 1 mm of media depth) provides improved spatial and temporal control over many culture parameters,^[13,14]^ allowing better mimicry of the dynamic and heterogeneous cellular microenvironment seen *in vivo*.^[15,16]^ For instance, *in vivo* the cell-to-cell, cell-to-environment interactions usually occur in a parenchyma (e.g. capillary bed) with a low volume ratio between interstitial fluid and cells, which defines the mass transport of various signalling factors (e.g. nutrients, vital gases, cytokines, etc.). Here we aim to develop a method that allows the cells to regulate the oxygen microenvironment and respond to changes autonomously at microscale, via a supply-demand balance as seen *in vivo*.

Theoretically, the oxygen microenvironment in microscale cell culture systems can be regulated in three ways based on the diffusion barrier set in between the supply and the demand: supply-dominated, demand-dominated, and supply-demand-balanced (Figure 1c-g).^[17]^ Various materials have been used to make microscale devices, including silicon, glass, plastic, and elastomer.^[18]^ Among them, polydimethylsiloxane (PDMS) has been most frequently used. While commonly referred to as PDMS, this elastomer is essentially a crosslinked, porous composite network of PDMS polymer/oligomer with various fillers/additives. Despite its wide acceptance, the ultra-high gas diffusivity of PDMS elastomer (**Table 1**) and the small scale of microscale cell culture systems [Figure 1c (left),d] render the oxygen microenvironment in PDMS-based systems supply-dominated. Oxygen levels in these systems are predominantly set and regulated by the supply, for instance by the ambient (i.e. the gaseous contents that the culture setup is exposed to) oxygen levels, via gas-regulation channels within PDMS devices, or by the continuous perfusion of oxygenated/deoxygenated media.^[17]^ Change from demand in the supply-dominated systems has limited influence on the oxygen microencrioment. The supply-dominated oxygen microenvironment imposes a preset value or gradient of oxygen level upon cells, which limits their spatiotemporal regulation and cellular response to the oxygen microenvironment in a physiologically relevant manner. By contrast, gas-impermeable plastics such as polystyrene (PS) and polymethyl methacrylate (PMMA) have been used to block oxygen diffusion into microscale cell culture chambers (Table 1).^[19]^ The oxygen microenvironment in these systems are demand-dominated [Figure 1c (right),e], i.e. predominantly defined by oxygen consumption of the cells within the system with limited supply regulation. Without continuous replenishment of oxygen supply (e.g. static media in sealed microwells),^[20]^ the demand-dominated oxygen microenvironment results in continuous decrease in oxygen levels to hypoxia and eventually to anoxia that leads to cell death. To mimic the kinetics and spatiotemporal variations in oxygen tensions characteristic of *in vivo* biology, cell culture systems should accommodate a Goldilocks or “just right” diffusion barrier for a supply-demand-balanced oxygen microenvironment as opposed to a strict supply-dominated or demand-dominated system. The supply-demand-balanced oxygen microenvironment allows continuous and dynamic regulation of the oxygen levels (i.e. POC and IOC) and cellular response via a supply-demand balance (Figure 1f,g).

**Table 1.** Reported oxygen solubility and diffusivity of materials typically used in microscale cell culture systems.

| Materials | Oxygen solubility<br>(ml of gas/1 l of fluid @ 1 bar of air) | Diffusion coefficient of oxygen<br>($\times 10^5$ cm<sup>2</sup>/s @ 1 bar of air) |
| --- | --- | --- |
| Deionized (DI) water | 5.4 (37 °C)<sup>[29]</sup> | 1.9-2.3 (25 °C)<sup>[30]</sup> |
| Typical cell culture media + 10% fetal bovine serum (FBS) | 5.3 (37 °C)<sup>[31]</sup> | 2.69 (37 °C)<sup>[31]</sup> |
| Silicone oil (i.e. SO) | 51.9 (5 cSt, 38 °C)<sup>[32]</sup><br>Data not found (1000 cSt) | 0.5 (500 cSt, 30 °C)<sup>[33]</sup><br>Data not found (5 cSt, 1000 cSt) (Figure S2) |
| Mineral oil | 49.5 (hexadecane, 22 °C)<sup>[32]</sup> | 2.49 (hexadecane, 22 °C)<sup>[32]</sup> |
| Fluorinert FC-40 (i.e. FC-40) | 76.8 (25 °C)<sup>[32]</sup> | 8.3 (22 °C)<sup>[32]</sup> |
| Polydimethylsiloxane (PDMS) elastomer | 310 (27 °C)<sup>[34]</sup> | 16 (27 °C)<sup>[34]</sup> |
| Polystyrene (PS)/Polypropylene (PP)/Polycarbonate (PC) | 174.7 (PS, 25 °C)<sup>[35]</sup> | 0.04-0.02 (25 °C)<sup>[36]</sup> |
| Polymethyl methacrylate (PMMA) | 194.8 (25 °C)<sup>[35]</sup> | 0.0025 (25 °C)<sup>[36]</sup> |

Recently, there has been renewed interest in the area of multiliquid-phase microfluidics, named under-oil open microfluidic systems (UOMS).^[21–28]^ In UOMS cell culture, aqueous media and cells are contained under an oil overlay, separating the cell culture microenvironment from the ambient with an immiscible liquid (i.e. oil) rather than solid materials used in traditional microscale devices (Figure 1f, Figure S1). Thus, compared to PDMS elastomer or other solid materials used in closed-channel or closed-chamber microscale devices, the oil overlay allows: (i) integration of a readily tailorable diffusion barrier for a supply-demand-balanced oxygen microenvironment by selecting/adjusting different oil properties (e.g. oil type, depth and viscosity), and (ii) facile and seamless intervention and spatially flexible deployment of external sensors (e.g. oxygen, pH, temperature, and etc.) on device to monitor the culture microenvironment in real time.

In this work, we demonstrate a method to establish and dynamically monitor AROM (i.e. autonomously regulated oxygen microenvironments) covering a full range of oxygen levels including hyperoxia, physioxia, hypoxia, and anoxia via a supply-demand balance in UOMS cell culture. Using a selected oil overlay, AROM can be achieved without the need of varying the media volume (i.e. the volume ratio between media and cells) and the media depth, which simultaneously would change cellular physiology and metabolism (e.g. nutrient availability, signaling efficiency). Moreover, AROM does not require media pre-deoxygenation (i.e. complete depletion of oxygen dissolved in the culture media) or external gas-regulation equipment, which streamlines the operation and lowers the adoption barrier among end users. We simulate the kinetics of oxygen diffusion against varying supply, demand, and diffusion barriers using COMSOL Multiphysics. Furthermore, a panel of cell types including various mammalian cells (epithelial, endothelial, stromal, immune), fungi and bacteria are examined for their capacity to regulate the oxygen microenvironment under oil. A key challenge is co-culturing oxygen-consuming human intestinal epithelium and anaerobic commensal bacteria that inhabit the gastrointestinal tract, due to their disparate oxygen demands. We apply the method to establish and characterize a co-culture of human primary intestinal epithelial cells and a highly prevalent human-associated intestinal species *Bacteroides uniformis* (*B. uniformis*) with these complex oxygen demands.

## 2. Results

### 2.1. COMSOL Multiphysics simulation of the kinetics of oxygen diffusion in multiliquid-phase microsystems

The kinetics of oxygen diffusion in a microsystem (flow-free) can be quantitatively described using Fick’s laws. Fick’s first law describes O_2_ flux (*J*) (i.e. mass transport of oxygen per unit time per unit area):

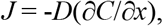

where

*D* is the diffusion coefficient of the diffusion barrier (e.g. ECM),

∂*x* is the distance between the supply and the cells,

-∂*C* is the difference in oxygen concentration between the supply and the cells, and

-∂*C*/∂*x* is the oxygen gradient (Figure 1a).

When O_2_ flux matches O_2_ demand, which can be defined by: O_2_ demand = oxygen consumption rate (OCR) (i.e. the amount of oxygen consumed per cell per unit time) × cell density (i.e. cells per unit area), the system reaches a supply-demand balance or oxygen homeostasis. The oxygen homeostasis can be shifted by changes from supply and/or demand (Figure 1g). For instance, higher oxygen supply results in increased POC and consequently IOC, whereas higher oxygen demand by cells results in decreased IOC and thereby a decrease in POC. Fick’s second law describes the change in oxygen concentration over time (∂*C*/∂*t*):

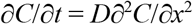

For a given ∂^2^*C*/∂*x*^2^, the inherently large gas diffusivity (i.e. *D*) in PDMS-based microdevices results in a large ∂*C*/∂*t*. Therefore, the large *D* and ∂*C*/∂*t* together result in the supply-dominated nature in the PDMS-based microdevices (i.e. high responsivity of POC and IOC to changes from supply), allowing little change in -∂*C*/∂*x* [the oxygen gradient established between the supply (i.e. either the ambient, gas-regulation channels, or the culture media) and the cells] when there is an altered O_2_ demand in the cells [Figure 1c (left),d]. Hence, in supply-dominated systems an altered O_2_ demand of the cells has little influence on POC and IOC, limiting relevance to the *in vivo* oxygen microenvironments where cells define the O_2_ demand, regulate the oxygen levels (i.e. POC and IOC), and respond to the oxygen microenvironment [e.g. via the hypoxia-inducible factor (HIF) signalling pathways]^[6,12]^ in a real-time feedback loop. By contrast, O_2_ flux is negligible (i.e. *J* → 0) in demand-dominated systems due to the limited gas diffusivity (i.e. *D* → 0) and thus oxygen in the media is continuously depleted by the cells and eventually leads to anoxia (i.e. POC and IOC = 0% O_2_) and cell death [Figure 1c (right),e].

On COMSOL Multiphysics (Experimental Section), we first investigated the influence of media depth on the kinetics of oxygen diffusion (**Figure 2**a,b). As expected, if the media depth is too small (< 1 mm), little decrease of oxygen level was obtained. By simply increasing the media volume and thus the depth, the full range of oxygen levels (i.e. hyperoxia, physioxia, hypoxia, and anoxia) can be achieved. However, increasing cell culture volume, thereby increasing the volume ratio between media and cells, changes the biology of the system confounding the data obtained. In comparison, an oil overlay instead provides an effective and flexible approach to achieve supply-demand-balanced oxygen microenvironments in microscale cell culture with a given volume ratio between media and cells.

**Figure 2.**
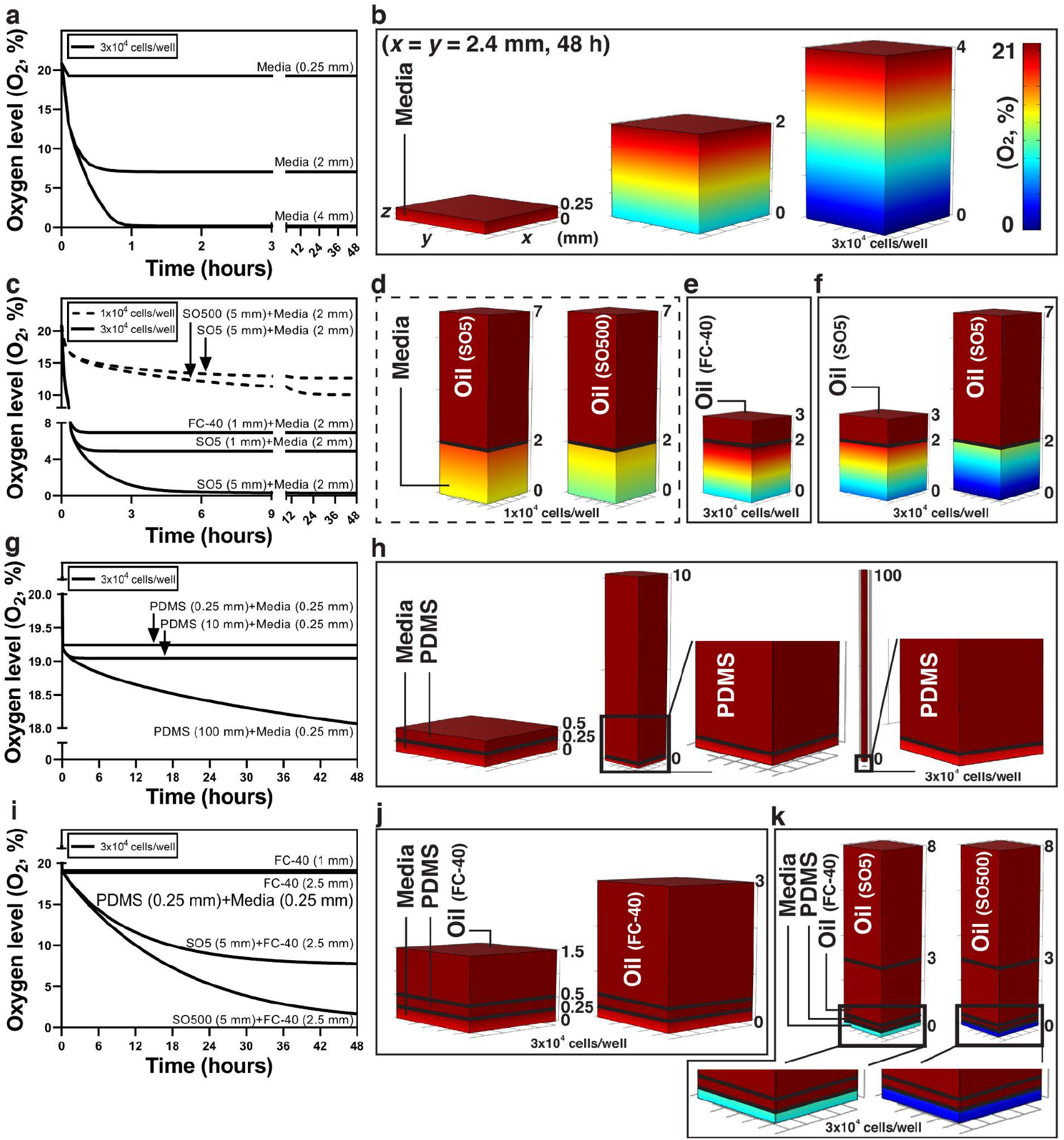
The kinetics of oxygen diffusion in multiliquid-phase microsystems and 3D diffusion profiles from COMSOL Multiphysics. a) and b) Oxygen level (O_2_, %) versus time plots and 3D diffusion profiles for the influence of media depth. c), d), e), and f) Oxygen level (O_2_, %) versus time plots and 3D diffusion profiles for the influence of oil type [fluorinated oil (FC-40) versus silicone oil (SO)], depth, and viscosity [5 cSt (SO5) and 500 cSt (SO500) of silicone oil]. g) and h) Oxygen level (O_2_, %) versus time plots and 3D diffusion profiles from PDMS (elastomer)-based microdevices. i), j), and k) Oxygen level (O_2_, %) versus time plots and 3D diffusion profiles from PDMS (elastomer)-based microdevices with oil (i.e. fluorinated oil) or double-oil (i.e. silicone oil + fluorinated oil) overlay.

In typical UOMS cell culture, the three most commonly used oil types are mineral (or paraffin) oil, silicone oil (i.e. pure linear PDMS polymer liquid), and fluorinated oil (i.e. perfluorocarbon liquid) due to their overall biocompatibility.^[37]^ However, in the context of gas permeability, the oxygen solubility and diffusivity of these oil types vary (Table 1). For instance, fluorinated oil shows notably high oxygen solubility and diffusivity and has been used as an oxygen carrier (i.e. to increase oxygen delivery) in cell culture.^[32]^ In comparison, oxygen solubility and diffusivity of silicone oil (which is viscosity-dependent) can be significantly lower than fluorinated oil. To control oxygen diffusion through the oil overlay we selected silicone oil as the gas diffusion barrier to get AROM in microscale cell culture. Moreover, silicone oil also allows Exclusive Liquid Repellency (ELR) - an extreme wettability in which a liquid gets absolutely repelled from a solid surface when exposed to a “right” secondary immiscible liquid. ELR enables versatile and advanced control of open fluids which has been reported in our previous UOMS publications.^[23,25,38]^

We performed the simulation with oil overlay against two cell seeding densities (i.e. 1×10^4^ cells/well versus 3×10^4^ cells/well), two oil types (i.e. fluorinated oil versus silicone oil), and two viscosities of silicone oil (i.e. 5 cSt versus 500 cSt) (Figure 2c-f). For a given volume ratio between media and cells higher viscosity of silicone oil leads to lower oxygen levels (Figure 2d), which is consistent with the oxygen diffusion test results (Figure S2). Due to the known high gas diffusivity, fluorinated oil doesn’t add much diffusion barrier to the culture media layer (Figure 2b,e). In comparison, silicone oil shows more significant influence on oxygen diffusion and thus the oxygen levels when applied on top of the culture medial layer (Figure 2f).

While not the focus of this work, we also investigated the oxygen diffusion through PDMS elastomer layers with small media depth (e.g. 0.25 mm) (Figure 2g-k). Not surprisingly, the PDMS elastomer layer shows little influence on oxygen diffusion until it reaches an awkwardly large thickness (Table 1, Figure 2h). Next, we simulated the function of oil overlay on top of the PDMS elastomer layer. It is worth noting that silicone oil causes severe swelling of PDMS-based devices, in contrast, fluorinated oil doesn’t cause any swelling issue on PDMS elastomer due to its chemical inertness. As expected, fluorinated oil doesn’t help much as a diffusion barrier (Figure 2j). Silicone oil plus fluorinated oil, the so-called “double-oil” overlay, especially with high viscosity of silicone oil, effectively affects the oxygen levels on PDMS-based microscale devices (Figure 2k). These simulation results were further confirmed in experiment (Figure S3).

### 2.2. Investigating possible oil extraction of lipophilic molecules from the culture media

A most frequently expressed concern during the development of UOMS cell culture was the possible extraction of lipophilic molecules by the oil phase from the culture media, which we also believe is an important attribute of the system that needs to be carefully inspected and understood before the use in cell cultures. The general biocompatibility of silicone oil [i.e. pure linear PDMS polymer liquid (without small molecule additives and cross-linkers)] in cell culture has been proven and reported. Here we adopt ultra performance liquid chromatography-tandem mass spectrometer (UPLC-MS) to systematically analyze the possible molecule loss in UOMS cell culture (**Figure 3**, Figure S4).

**Figure 3.**
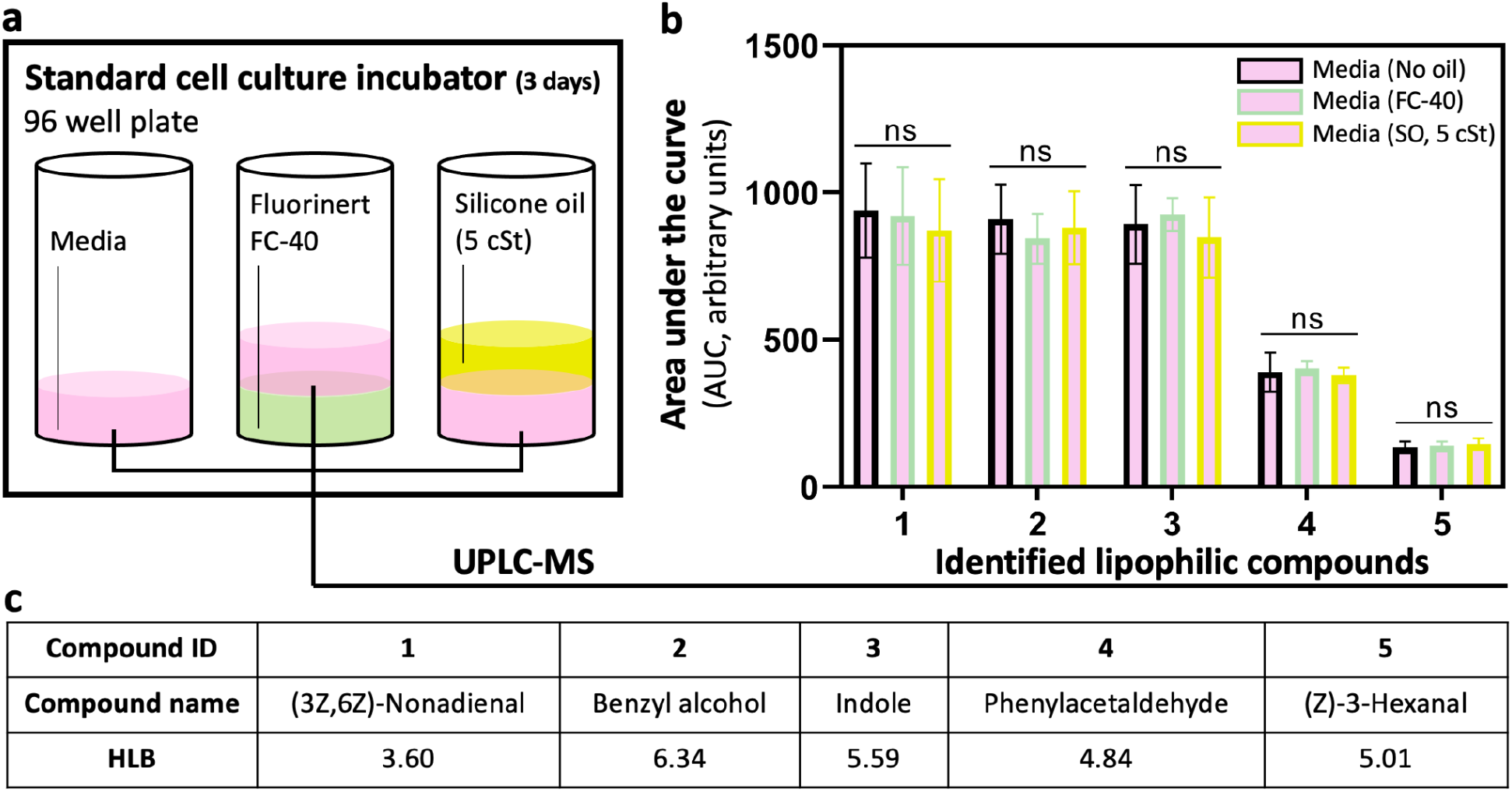
UPLC-MS media analysis for possible oil extraction of lipophilic molecules. a) Schematic shows the test conditions. The culture media was incubated for 3 days without agitation before collection and analysis. b) Comparison of the relative abundance (i.e. area under the curve, or AUC from the extracted ion chromatograms, or EICs) (Figure S4) of five identified lipophilic compounds. “ns” represents “not significant”. c) The compound names in (b) with the HLB values.

We run the test with a typical culture media [Dulbecco’s Modified Eagle’s medium (DMEM) + 10% FBS + 1% Pen-Strep] on a standard polystyrene 96 well plate in a standard cell culture environment, i.e. 37 ^°^C, 18.6% O_2_, 5% CO_2_, and 95% relative humidity (RH) (Figure 3a, Experimental Section). The media was collected and analyzed on UPLC-MS after 3 days of incubation without agitating the oil and media layers. The identified molecules from UPLC-MS were ranked by their hydrophilic-lipophilic balance (HLB) values. We screened out the top five lipophilic molecules with HLB ranging from 0 to about 6 (Figure S4) for comparison between the no oil and oil conditions (Figure 3b,c). Not surprisingly, fluorinated oil shows no molecule loss compared to no oil due to its widely reported chemical inertness. Interestingly, we didn’t observe any significant molecule loss from silicone oil either even against the lipophilic molecules. Actually, the retention of small molecules in water-in-oil emulsion droplets has been broadly studied in droplet microfluidics.^[39]^ In general, the molecule retention or loss highly depends on the lipophilicity of the target molecules and the surfactant layer (i.e. surfactant type and concentration) at the oil-media interface (Figure S5). It has been reported the addition of biopolymers (e.g. bovine serum albumin, or BSA) significantly improves the retention of lipophilic molecules in the media phase.^[40]^ The high retention of lipophilic molecules in the culture media from the UOMS cell culture streamlines the parameter space when used in cell culture and cell-based assays.

### 2.3. Kinetics of POC and IOC in UOMS cell culture with varying supply-demand balances

The oxygen microenvironment *in vivo* is highly heterogeneous across organs.^[1]^ In local tissue, the POC and IOC are also varied spatially per the distance between the cells and the supply (i.e. capillaries), which are delicately defined by the supply-demand balance via oxygen diffusion. For example, a steep oxygen gradient is known to exist in the intestine with 4-8% O_2_ in the submucosa and lamina propria, 2-4% across the epithelial and mucous layer, and less than 2% in the lumen.^[7]^ We chose a representative cell line of the intestinal tissue (human colorectal tumor cell line, Caco-2) to demonstrate the capability to recapitulate different local oxygen levels of tissue *in vivo* via AROM in UOMS cell culture (**Figure 4**a,b). The POC in UOMS cell culture can be monitored at any time point by directly introducing single or multiple external oxygen microsensors through the oil overlay. To facilitate dynamic monitoring of the kinetics of oxygen levels with single-cell resolution, we used a hypoxia dye taken up by live cells to probe the IOC in parallel with the POC measurement. The combined POC and IOC measurements (Experimental Section) allow a reliable and multifaceted way of monitoring the oxygen microenvironment in UOMS cell culture.

**Figure 4.**
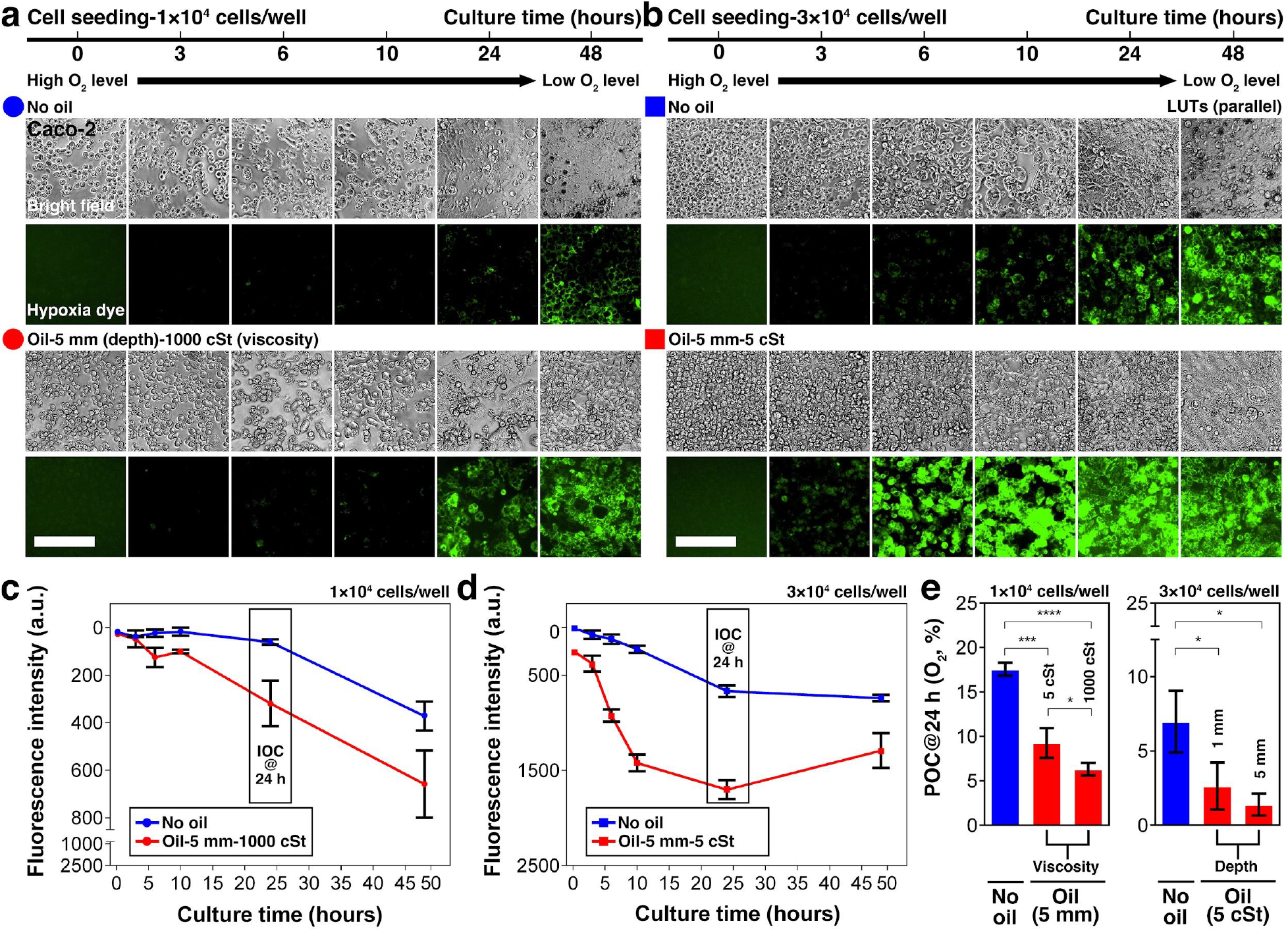
Kinetics of POC and IOC in UOMS cell culture with different seeding densities of cells and selected oil properties. An intestinal epithelium cell line (Caco-2) was cultured in two different seeding densities with and without oil overlay: a) 1×10^4^ cells/well, and b) 3×10^4^ cells/well. 3×10^4^ cells/well leads to a near-confluent monolayer from cell seeding in a standard 384 well. The volume of media was set constant at 20 μl/well (for 2 mm in media depth). 10 or 50 μl/well of silicone oil (5 cSt or 1000 cSt) was added on top of the media layer (for an additional 1 or 5 mm in oil depth). The fluorescent images of hypoxia dye were processed with parallel lookup tables (LUTs) (i.e. the same minimum and maximum in brightness/contrast adjustment) for the comparison of fluorescence intensity. See Figure S7 for large images of the typical cell morphologies. Scale bars, 200 μm. c) and d) The kinetics of IOC (monitored by the hypoxia dye) in 48 h corresponding to (a) and (b), respectively. e) The influence of oil viscosities (5 cSt versus 1000 cSt) or depths (1 mm versus 5 mm) shown by POC (O_2_, %) measured at 24 h using the optical oxygen sensor (Figure S1). Data were pooled and averaged with ×3 replicates in each condition. Error bars, mean ± s.d. \**P* ≤ 0.05, \*\**P* ≤ 0.01, \*\*\**P* ≤ 0.001, and \*\*\*\**P* ≤ 0.0001.

Here, we explore the simulated kinetics of oxygen diffusion after cell seeding (Figure 2), and examine in experiment the influence of cell seeding density and oil properties (i.e. oil depth and viscosity) on POC and IOC.

In this study, cell culture media was normally saturated with oxygen from exposure to atmospheric concentration (i.e. 21% O_2_). Immediately after cell seeding, no oxygen gradient has been established yet (i.e. -∂*C*/∂*x* = 0) and thus, O_2_ flux is zero (i.e. *J* = 0). Over time, cells consume the dissolved oxygen from the surrounding media, thereby establishing an oxygen gradient that increases the difference between ambient and the local media (i.e. -∂*C*/∂*x* > 0), leading to an increased O_2_ flux (i.e. *J* > 0). Eventually, O_2_ flux reaches its maximum, either determined by (and equal to) O_2_ demand or less than the demand, at which point diffusion becomes the rate-limiting step. In the case of O_2_ flux matching O_2_ demand, POC and IOC reach a steady state (or homeostasis). In the case of O_2_ flux less than O_2_ demand (i.e. inadequate oxygen delivery to the cells), POC and IOC drop to 0% O_2_ which leads to anoxia. The discussion above assumes a constant O_2_ demand. Note that O_2_ demand may vary over time due to cell proliferation, cell death, or changes in metabolic activities. To achieve a quasi-constant O_2_ demand (e.g. within 24 h of culture), we chose the seeding density of 3×10^4^ cells/well which produces a near-confluent monolayer from cell seeding in a standard 384 well. In such conditions, cell proliferation is not a dominant parameter that significantly varies O_2_ demand considering contact inhibition and limited surface area. To investigate the influence of cell growth on the kinetics of AROM, we used a lower seeding density (1×10^4^ cells/well) in parallel as comparison. In all tested conditions, cell viability was maintained at a normal level (Figure S6). Note that the conditions with lower cell-media ratio and/or oil overlay exhibited higher cell viability.

Here, we demonstrate that our experimental results captured the simulated kinetics of oxygen diffusion as described above. We summarize several guidelines to control the kinetics of the AROM in UOMS cell culture:

i. Higher cell seeding density or larger oil depth/viscosity leads to lower POC and IOC (Figure 4c-e).
ii. For the default cell seeding density (i.e. 3×10^4^ cells/well in a standard 384 well) and cell type (i.e. Caco-2), POC and IOC reach a steady state (or homeostasis) in around 24 h from the initiation of culture (Figure 4d). It takes a longer time for the oxygen microenvironment to reach homeostasis with lower cell seeding density or a cell type with lower OCR (Figure 4c).
iii. Varying the oil depth in the cultures with a confluent monolayer of Caco-2 cells (the 3×10^4^ cells/well condition) created oxygen levels representative of different regions of the intestine: the submucosa (no oil overlay, ∼7% O_2_), epithelial and mucous layer (1 mm oil overlay, ∼3% O_2_), and lumen (5 mm oil overlay, < 2% O_2_) (Figure 4e, right).
iv. Different oxygen levels (Figure 1a) can be obtained without using oil overlay, for example, by only changing the media depth^[31]^ (Figure 2a, Figure 4e, Figure S8). However, as aforementioned, the change of media volume in a culture also changes many other factors such as nutrient availability and the concentration of metabolites or signaling molecules that together compound the readouts.

Taken together, the results validate the high flexibility and controllability of establishing AROM in UOMS cell culture with a simple oil overlay.

### 2.4. Cellular regulation of the under-oil oxygen microenvironment

A typical *in vivo* cellular microenvironment includes multiple cell types, and in many cases inter-kingdom interactions. It is well known that a broad heterogeneity of OCR exists across cell types.^[31]^ Here, a panel of mammalian and fungal cell types (Figure S9, Table S1) are examined for their capacity to regulate the oxygen microenvironment in UOMS cell culture. This test also demonstrates the capability to perform compartmentalized and high-throughput oxygen regulation on one device with a single, shared ambient environment (**Figure 5**a).

**Figure 5.**
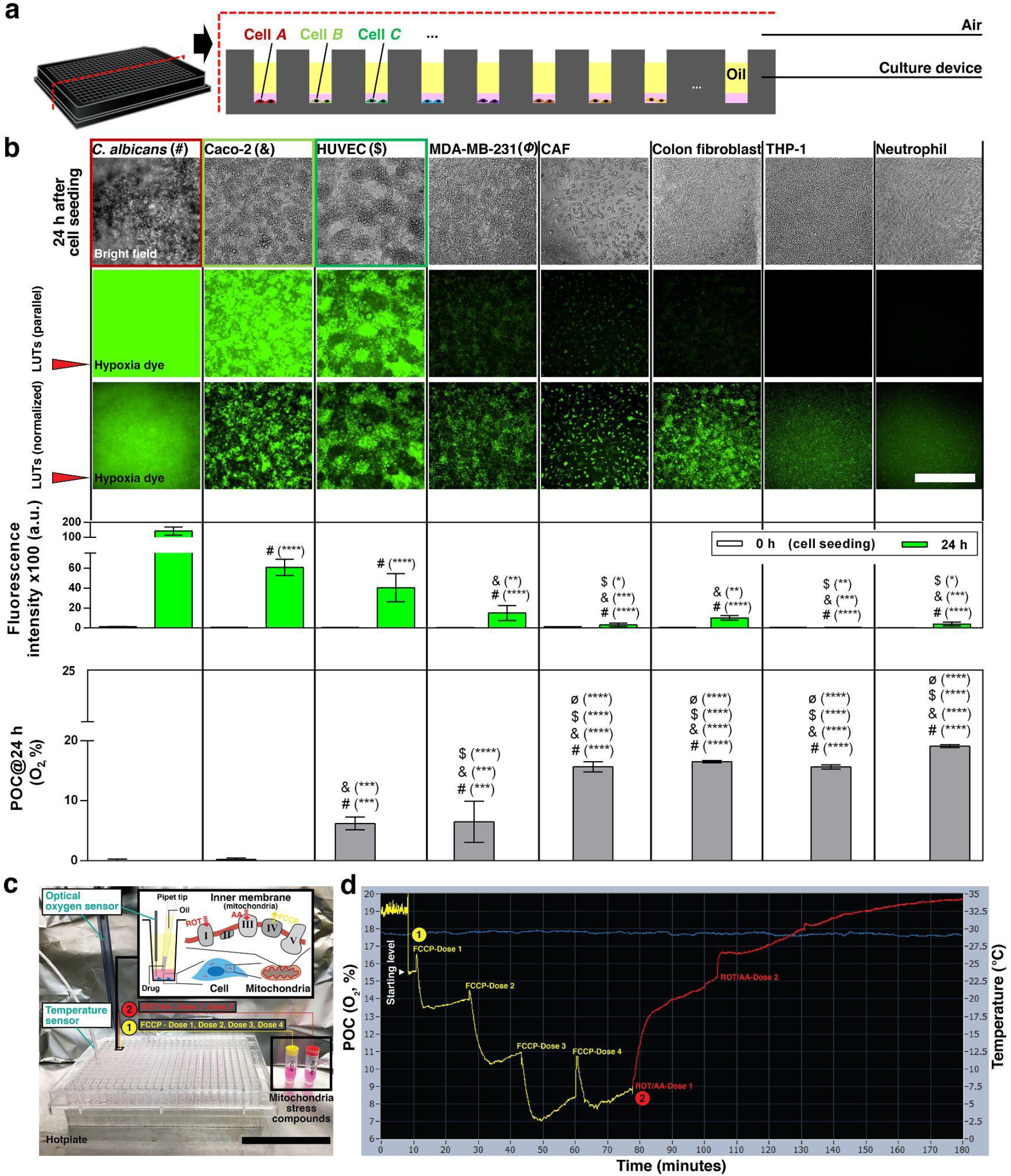
Under-oil oxygen microenvironments regulated by different cell types. a) Schematic shows the compartmentalized, high-throughput oxygen regulation on one device (e.g. a 384-well plate) with different cell types and a shared atmosphere (e.g. air). b) Each cell type was seeded at 3×10^4^ cells/well on a 384-well plate with 20 μl/well of media (for 2 mm in media depth), overlaid with 50 μl/well of silicone oil (5 cSt) (for 5 mm in oil depth), and cultured up to 24 h for a parallel comparison. The fluorescent images of hypoxia dye were processed with parallel LUTs (the middle row) for the comparison of fluorescence intensity, and with normalized LUTs (i.e. individually adjusted minimum and maximum in brightness/contrast) (the bottom row) for visualization. See Figure S10 for large images of the typical cell morphologies. Scale bar, 500 μm. The bar graph of POC (O_2_, %) and IOC (fluorescence intensity of hypoxia dye) of each cell type was displayed under relevant panels of the fluorescent images. Data were pooled and averaged with ×2 cell passages and a minimum of ×3 replicates of each condition. Error bars, mean ± s.d. \**P* ≤ 0.05, \*\**P* ≤ 0.01, \*\*\**P* ≤ 0.001, and \*\*\*\**P* ≤ 0.0001. c) Reconditioning of under-oil oxygen microenvironments and the experimental setup for regulation of mitochondrial respiration in UOMS cell culture. Caco-2 cells [1×10^4^ cells/well, 20 μl/well of media (for 2 mm media depth), 50 μl/well silicone oil (5 cSt) (for 5 mm oil depth) overlay] were cultured for 12 h to reach a starting level of POC at about 15% O_2_. Two sets of mitochondrial stress compound [carbonyl cyanide-4 (trifluoromethoxy) phenylhydrazone (FCCP, an uncoupling agent that maximizes the oxygen consumption of Electron Transport Chain (ETC) Complex IV), and a mixture of rotenone (ROT, an inhibitor of ETC Complex I) and antimycin A (AA, an inhibitor of ETC Complex III)] were added to a culture well (0.4 μl/dose) in a sequence to manipulate the mitochondrial respiration of the cells (Experimental Section). The pipette tip for compound delivery was prefilled with oil to avoid introducing air to the media under oil during pipetting. Scale bar, 5 cm. d) Real-time (3 h) POC (the yellow-red line) of the reconditioning with regulated mitochondrial respiration. The oxygen sensor was set on the cell layer without being moved during the entire measurement process. The O_2_ peak immediately after each compound loading was attributed to the dissolved oxygen in each dose (0.4 μl) of the compound solution. The oxygen level of media (no cells) at the test temperature (∼30 ^°^C, the blue line) is about 20% O_2_.

Within the cohort of mammalian cells, cancer cell lines [Caco-2 (a human colorectal adenocarcinoma),^[41]^ MDA-MB-231 (a migrating breast cancer)^[42]^], and endothelial cells [HUVEC (human umbilical vein endothelial cell)^[43]^] show high OCR and thus high capacity of oxygen regulation according to both POC and IOC measurements (Figure 5b). In comparison, fibroblasts [CAF (cancer-associated fibroblasts, breast cancer tissue), colon fibroblasts (normal tissue)] and the two types of non-adherent white blood cells [THP-1 (a human monocytic cell line), neutrophils (primary)^[44]^] show a relatively lower OCR and capacity of oxygen regulation compared to other cell types. The fungus (*Candida albicans*, or *C. albicans*) dwarfs all of the mammalian cells with respect to a drop in POC and IOC levels due to its rapid growth rate and high OCR^[45]^ relative to mammalian cells. It’s worth noting that the POC and IOC levels for each cell type examined in UOMS cell culture are consistent with reported OCRs and the typical metabolic pathways taken by those cell types.^[41–45]^ Indeed, in a cellular microenvironment involving multiple cell types, the OCR of individual cell types may change via cell-to-cell signaling or cross-talks. Systematically examining the influence of the signaling factors on OCR in combinatorial co-cultures is beyond the scope of this investigation. The results in this section provide a basis for the capacity of oxygen regulation of each cell type prior to establishing inter-kingdom co-culture systems with AROM.

*In vivo*, the oxygen microenvironment is dynamically regulated and reconditioned as the supply-demand balance shifts in response to various stressors. Here, we seek to demonstrate reconditioning of the AROM in UOMS cell culture that mimics the *in vivo* oxygen homeostasis processes. We adopted a standard mitochondrial stress assay kit to manipulate the mitochondrial respiration and thus the OCR (or O_2_ demand) of the cells (Experimental Section) (Figure 5c). The POC of the UOMS cell culture was monitored in real time for 3 h (Figure 5d). The results showed a fast and robust response of the AROM to the mitochondrial stress compounds with a series of reconditioned oxygen levels (i.e. the characteristic supply-demand-balanced oxygen response and homeostasis shown in Figure 1g) successfully captured.

### 2.5. Investigating a co-culture between primary colon epithelial cells and human-associated anaerobic intestinal bacteria using UOMS cell culture

A major challenge to studying biological systems is the development of co-culture models that can recapitulate key parameters of natural microenvironments.^[46]^ In this section, we explore an *in vitro* co-culture between intestinal epithelium and anaerobic bacteria, which presents unique challenges due to their distinct demands for and tolerance to oxygen. A specific oxygen microenvironment must be established to sustain metabolic activities of host cells and bacteria with physiological or pathological relevance. While several *in vitro* co-culture models have been developed to study host-microbe interactions utilizing microscale cell culture,^[47,48]^ the oxygen levels in these systems are typically supply-dominated (or demand-dominated) as previously discussed, imposing a physiologically inconsistent oxygen microenvironment to the cells. In addition, the adoption barrier of these *in vitro* models, including limited access to samples within the devices due to the closed-channel or closed-chamber designs as well as the access to specific equipment for device fabrication and operation, have stymied their broader adoption. Here, we apply the UOMS cell culture with AROM to establish a co-culture between primary colon epithelial cells and human-associated anaerobic intestinal bacteria. We characterize and validate the co-culture using measurements of POC, gene and protein level expression of differentiation markers of the intestinal epithelium, and bacterial growth.

Although investigators have traditionally relied on colon cancer cell lines to study gut epithelial physiology, they may possess non-physiologic characteristics including altered metabolism, and aberrant proliferative and differentiation characteristics which call into question their predictive ability in modeling normal epithelial function.^[49,50]^ In this co-culture experiment, we cultured monolayers of colon epithelium derived from primary tissue on a thin layer of ECM coated culture plate (**Figure 6**a, Experimental Section) to represent a more physiologically relevant cellular microenvironment. Colon monolayers were overlaid with oil for 24 h to establish AROM prior to inoculation with anaerobic bacteria. Immunofluorescence staining (IFS) of E-cadherin and ZO-1 confirmed formation of epithelial adherens junction and z-stack modeling was used to determine monolayer formation (Figure 6b, Figure S11, Table S2) in the UOMS cell culture. In the gut, intestinal stem cells are able to differentiate into specialized cell types including absorptive enterocytes and mucus-producing goblet cells which make up the intestinal epithelium separating the gut lumen from the underlying parenchyma.^[51]^ In our UOMS cell culture, IFS of colon monolayers further confirmed expression of cell lineage markers such as mucin 2 (goblet cells), FABP1 (enterocyte) and villin (enterocytes expressing microvillar actin-binding protein) (Figure S11), indicative of a differentiated epithelium. Qualitative assessment of the IFS data showed no significant differences between the control (i.e. no oil overlay) and the AROM conditions (i.e. oil overlay) indicating biocompatibility of the oil overlay (silicone oil) in the *in vitro* culture of primary cells.

**Figure 6.**
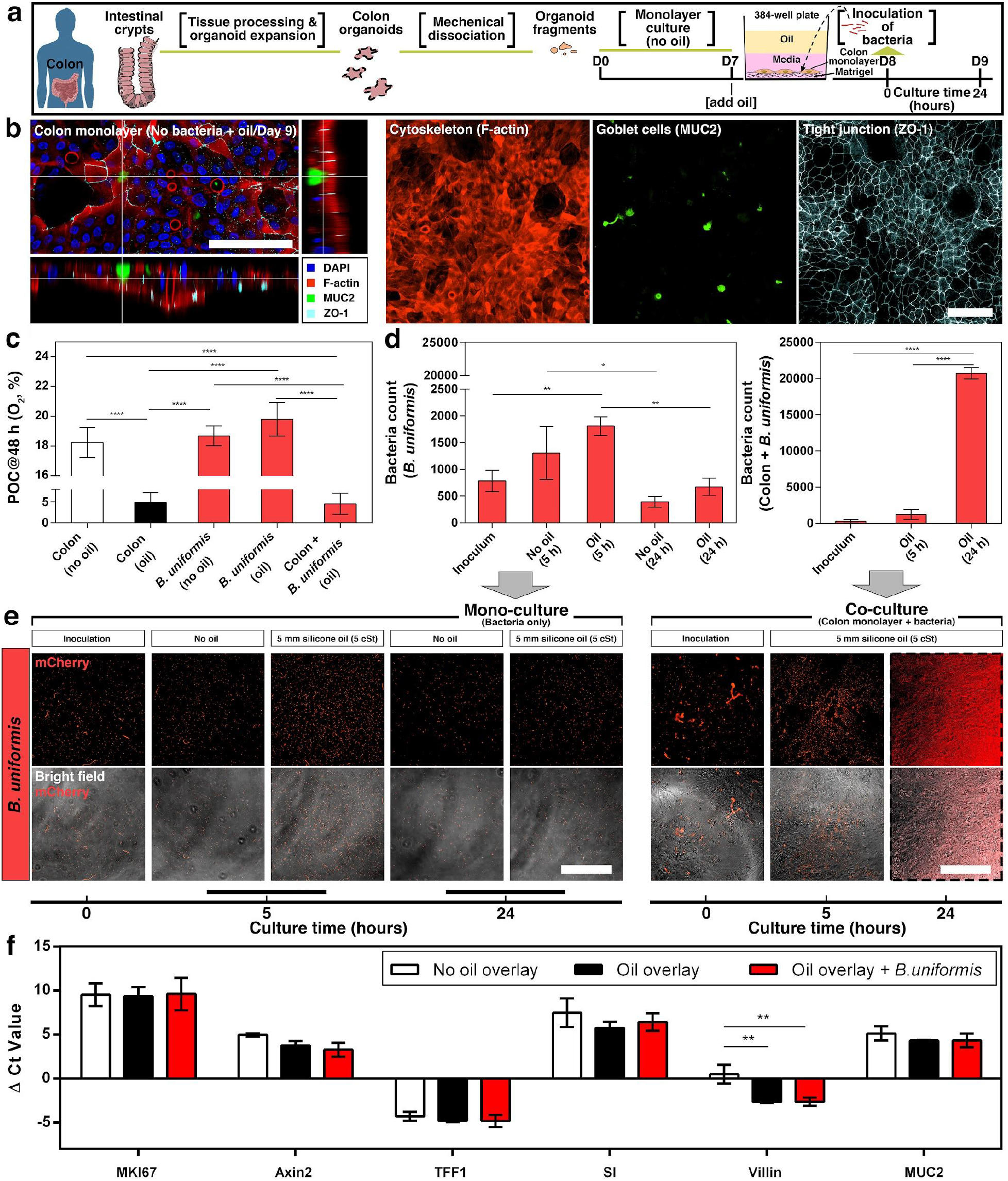
Under-oil co-culture, cell function of primary colon epithelium, and growth kinetics of anaerobic bacteria (*B. uniformis*). a) Schematic shows the workflow of tissue processing, organoid generation, establishment of colon monolayer, and inoculation of bacteria. The volume of media is 40 μl/well for 4 mm in media depth. The volume of oil (silicone oil, 5 cSt) is 50 μl/well for 5 mm in oil depth. b) Three-dimensional (3D) laser confocal image with z-slices and fluorescent images from the under-oil mono-culture of colon monolayer on Day 9. Scale bars, 100 μm. c) POC (O_2_, %) measured on Day 9 (i.e. at 48 h from adding the oil overlay) along with two no-bacteria mono-culture conditions (i.e. colon monolayer only with and without oil overlay). Data were pooled and averaged with a minimum of ×3 replicates of each condition. d) Bacteria count [at 30× magnification, with a field of view of 446.5 μm (length) × 445.6 μm (width)] in (e). e) Microscopic images (30× magnification) showing the growth in mono-culture (bacteria only) with and without oil overlay compared to co-culture (colon monolayer + bacteria) under oil. The fluorescent images were all processed with normalized LUTs for visualization. The 24 h *B. uniformis* images (in the dashed line box) were rendered with pseudocolor (red) to visualize the bacteria (Experimental Section). Scale bars, 200 μm. f) RT-qPCR [ΔCt (i.e. cycle threshold)] results for comparison of gene expression (Table S2) of the colon epithelium. Lower ΔCt values indicate higher gene expression. Data were pooled and averaged with ×3 replicates of each condition. Error bars, mean ± s.d. \**P* ≤ 0.05, \*\**P* ≤ 0.01, \*\*\**P* ≤ 0.001, and \*\*\*\**P* ≤ 0.0001.

The adult intestinal microbiota consists of hundreds of species, majority of which are obligate anaerobes, which are inhibited by the presence of oxygen due to their inability to defend against cellular damage by reactive oxygen molecules.^[52]^ In the colon, oxygen is consumed by luminal contents and the resident tissue cells to maintain the growth of strict anaerobes.^[7]^ We hypothesized that by leveraging AROM by primary intestinal epithelial cells in UOMS cell culture, the oxygen level can be reduced to the physiological level that facilitates co-culture and growth of strict anaerobes such as *Bacteroides. Bacteroides* are a highly prevalent and abundant group of species in the human gut microbiome that can benefit the host by performing chemical transformations that provide key nutrients to the host.^[53]^ In general, *Bacteroides* are strict anaerobes but specific species such as *B. fragilis* have been shown to tolerate and benefit from nanomolar oxygen.^[54]^ Here, we used *B. uniformis*, genetically modified to constitutively express mCherry fluorescent protein (mCherry) to demonstrate the utility of our method in facilitating co-culture of primary intestinal cells with human-associated anaerobic bacteria. POC measurements of the colon monolayer [Figure 6c, Colon (no oil/oil)] showed that with the oil (silicone oil, 5 cSt) overlay the oxygen level can be autonomously reduced to 2-8%, consistent with the physioxia of intestinal tissue, compared to no-oil controls where oxygen levels remained at hyperoxia (17-19% O_2_). To investigate the OCR of *B. uniformis*, POC measurements were performed of *B. uniformis* mono-cultures with and without oil overlay. In spite of the oil overlay, the oxygen levels were maintained close to atmospheric 21% O_2_ over a 24 h period indicating a negligible consumption of oxygen by *B. uniformis* [Figure 6c, *B. uniformis* (no oil/oil)]. Further substantiating this observation, the co-culture of *B. uniformis* with colon monolayers yielded similar POC levels (2-8% O_2_) to colon cell mono-cultures indicating that oxygen consumption was dominated by colon epithelium.

We investigated the growth kinetics of *B. uniformis* in the mono-culture and co-culture conditions by quantifying the number of bacterial cells using fluorescent microscopy (Figure 6d,e). In bacterial mono-cultures, *B. uniformis* showed a moderate increase in abundance (Experimental Section) at an earlier time point of ∼5 h and then a decrease in abundance by 24 h. In contrast to the mono-culture condition, *B. uniformis* exhibited reciprocal growth trends in the presence of colon monolayers in UOMS cell culture. Our results indicate that the growth of *B. uniformis* was significantly enhanced in the presence of the intestinal cells, leading to a dense and confluent layer blanketing the colon monolayer. One potential explanation for the observed growth enhancement is the reduced oxygen levels due to the metabolism of the intestinal epithelial cells under oil. Corroborating these trends, the oxygen microenvironments were substantially different in the mono-culture and co-culture conditions at 48 h based on the POC measurements [Figure 6c, *B. uniformis* (no oil/oil) versus Colon + *B. uniformis* (oil)]. To further characterize the effect of oxygen on the growth of *B. uniformis* in the co-culture, we performed a co-culture experiment under fluorinated oil (Figure S12). Compared to silicone oil, fluorinated oil has low moisture and high gas (e.g. O_2_, CO_2_) permeability (Figure 2a-f, Table 1). As shown in Figure S12, little hypoxia was generated from the co-culture of the host cells (Caco-2) and the bacteria under fluorinated oil compared to co-cultures under silicone oil. Importantly, no growth of *B. uniformis* was observed in the co-culture performed under fluorinated oil in contrast to the growth observed in the co-culture under silicone oil. While factors beyond oxygen may affect the growth of *B. uniformis*, the fluorinated oil control experiment suggests that elevated oxygen may contribute to the impaired growth of *B. uniformis* in this co-culture relative to the silicone oil overlay.

The IFS analysis showed that expression of epithelial differentiation markers in colon monolayers were similar in presence and absence of *B. uniformis* (Figure S11), suggesting that the presence of *B. uniformis* did not significantly disrupt the epithelium. To investigate the impact of *B. uniformis* and UOMS cell culture on colon monolayers at the transcriptional level, we examined the expression of gene markers associated with epithelial cell function, including cell proliferation (MKI67) and cell lineage (Axin2, TFF1, SI and MUC2) (Table S2). Our results showed that all investigated genes were robustly detected in all conditions (no oil overlay, oil overlay and *B. uniformis* co-culture with oil overlay) by quantitative reverse transcription polymerase chain reaction (RT-qPCR) (Experimental Section). In addition, there were no significant changes in the expression of the proliferation marker gene (MKI67) and majority of characteristic cell lineage markers associated with differentiation (Axin2, TFF1, SI and MUC2) in response to UOMS cell culture (Figure 6f). Notably, high abundance of *B. uniformis* did not significantly alter the differentiation status of the epithelium according to the subset of genes examined, which is consistent with the IFS analysis results. However, our results indicate that the colon monolayer expression of villin, associated with brush border microvilli expressed by absorptive enterocytes, was upregulated in the conditions with UOMS cell culture compared to the control (i.e. no oil overlay) condition. Future studies are required to determine if the UOMS cell culture facilitates microvilli differentiation by upregulating the expression of villin and to confirm microvilli formation. Together, these results lend support to the hypothesis that the reduced oxygen level enabled by AROM in UOMS cell culture can establish a more favorable environment for the growth of strict anaerobes such as *B. uniformis*.

## 3. Discussion

The critical role of oxygen in living systems is increasingly understood, however, precise monitoring and control of oxygen levels are often neglected in cell culture studies. Many studies have highlighted the importance of mimicking *in vivo* physiological conditions (e.g. nutrients availability, temperature, pH and the partial pressure of vital gases such as oxygen and carbon dioxide of the cellular microenvironment), to improve modeling of specific cellular microenvironments in human physiology and disease. Nevertheless, employing a physiologically inconsistent oxygen microenvironment in cell culture is still a widely existing pitfall, which likely contributes substantially to differences between *in vivo* and *in vitro* results.^[15]^ While the development of microscale cell culture enables precise control of the cellular microenvironment compared to standard cell culture at bulk scale, oxygen microenvironments that allow cellular control of the oxygen levels and achieve homeostasis (i.e. supply-demand balance) are lacking due to the material properties and small scale nature of microscale technologies. In this work, we introduce UOMS cell culture, which provides a reliable approach capable of recapitulating a full range of oxygen levels via autonomous regulation by cells. Oxygen regulation by cells allows for establishment of feedback loops wherein the dynamically changing oxygen demand of cells modifies the oxygen microenvironment, which in turn triggers signaling pathways that affect cell functions and phenotypes. Using our method, the oxygen microenvironment is not imposed on the cells by the operators and is thus distinct from existing microscale cell culture methods (which primarily employ a supply-dominated or demand-dominated gas control on the cell culture system). By contrast, the cells are allowed to control oxygen autonomously via a supply-demand balance, which resembles the regulation of the oxygen microenvironment *in vivo*.

While this method aims to establish autonomous regulation of oxygen levels for *in vitro* microscale cell culture, it could be applied to other vital gases (e.g. carbon dioxide) in UOMS cell culture (Figure S13). For microscale cell culture, especially PDMS-based microdevices, the dissolved gases in the culture media can rapidly change (due to the ultra-high gas permeability of PDMS elastomer and the small scale of the microdevice) if the ambient gas compositions change (e.g. moving the setup out of the incubator for intermediate sample manipulation or imaging). While this issue can be mitigated using a gas-regulation chamber equipped with a glove box and other tools (e.g. liquid handler, microscope), the complexity of such gas-regulation systems burdens the operation and discourages end users from adopting those measures. UOMS cell culture minimizes the dependence on a specifically defined ambient for vital gas regulation. A culture or co-culture can be established, maintained, and characterized directly in an atmospheric environment without using gas-regulation equipment.

To make the autonomous gas regulation complete as a scientific tool (i.e. being able to establish a specific microenvironment and perform quantitative measurements for tracking and investigating the process) the local level of such vital gases in the culture media to which the cells are exposed need to be effectively monitored. Compared to the culture systems with closed-channel or closed-chamber designs, UOMS cell culture allows minimal disturbance of the established cellular microenvironment from external interventions. As demonstrated in this work, a microsensor can be introduced through the oil overlay with spatiotemporal flexibility to monitor the local oxygen levels at sites of interest. Similarly, cellular samples and/or reagents (e.g. drugs, conditioned media) can be easily added to or collected from UOMS cell culture without interfering with the gas levels in the cellular microenvironment. Moreover, by arranging different cell types and modulating the oil type, depth or viscosity of the oil overlay, compartmentalized and high-throughput gas regulation can be realized on one device (e.g. a microtiter plate) with the same ambient environment such as in air or in a standard incubator. Temporarily changing oxygen levels seen *in vivo* such as fluctuating or cycling hypoxia can be also easily implemented by adding and removing a specific volume of oil in the oil overlay (e.g. with a programmed syringe pump) over time. The features enabled by the under-oil open design, could allow AROM to be used for high-throughput screening applications (e.g. mutant analysis of bacterial species or antimicrobial drug selection) with individually controlled oxygen microenvironments.

For *in vitro* microscale cell cultures that need to be maintained from several days to weeks, nutrients need to be replenished via media changes. To minimize interruption to the established oxygen microenvironment in UOMS cell culture during media change, we have demonstrated the use of partially deoxygenated media described in Figure S2. Deoxygenation of the culture media can be executed in a number of different ways including nitrogen gas (N_2_) bubbling, or a regular degassing/gas-exchange vacuum desiccator. The deoxygenated media can subsequently be stored under oil, and monitored using the oxygen sensor system as the oxygen level recovers to the target value (i.e. the oxygen level in an established microenvironment) required for media change.

Recently we reported a breakthrough in UOMS:^[38]^ Enabled by the extreme wettability - Exclusive Liquid Repellency (ELR, an inherent and absolute repellency of a liquid on a solid surface),^[23,25]^ open microchannels can be prepared under oil (silicone oil) with the channel dimensions reduced up to three orders of magnitude (from millimeter scale to micrometer scale) compared to previously reported techniques. Open-fluid cell trapping (including mammalian cells and bacteria), flow rate range comparable to blood flow with open-channel designs, and anti-biofouling reversible open-channel fluidic valves were achieved in open microfluidics. Designer microchambers/microchannels and versatile fluidic control can be easily and robustly introduced to UOMS cell culture with these recent advances from our lab. One of the ongoing follow-up works is to extend the AROM method to the under-oil open microchannels that allow various flow conditions.

We used the AROM method to investigate inter-kingdom interactions between primary host intestinal epithelium and a highly prevalent human gut microbiome species which exhibit contrasting responses to oxygen. Our results demonstrated that UOMS cell culture with AROM can be used to study the dynamics of inter-kingdom interactions at multiple levels including bacterial growth, host-cell phenotypes and gene expression. Future work will investigate the molecular basis of interactions using metabolomics, as well as study interactions using a diverse panel of human gut bacterial species and microbial communities constructed from the bottom-up. In addition, future studies could investigate the genetic determinants of the bacterial growth dynamics in the presence and absence of host epithelium in AROM using barcoded transposon libraries^[55]^ and/or deletions of key genes involved in oxygen sensing or tolerance.^[54,56]^ Further, the ecological and molecular role of oxygen as a mediating factor in gut microbiome dysbiosis could be investigated using AROM. Specifically, shifts in microbial community composition, metabolite production and degradation, and host phenotypes could be quantified in response to oxygen perturbation in order to elucidate disease relevant host-microbiome interactions and feedbacks.^[57]^ In addition, we will investigate the integration of 3D organotypic model components (e.g. incorporation of crypt/villus architecture, immune cells, vasculatures and media circulation) with the under-oil oxygen microenvironments.

The ability to control physical and chemical characteristics of cell culture in *in vitro* modeling will enable improved function (and relevance) when recapitulating normal and disease states seen *in vivo*. We foresee many potential applications of AROM in UOMS cell culture as providing improved capacity to mimic *in vivo* conditions, and as a functionality module that can be robustly and readily integrated into existing *in vitro* microscale cell culture systems allowing for broad adoption among end users.

## 4. Experimental Section

### COMSOL Multiphysics simulation

We performed the simulation on COMSOL Multiphysics (Ver. 5.6) with the Chemical Reaction Engineering (CRE) Module. We set temperature at 310.15 K and pressure at 1 atm as the default model inputs. We established the models based on the geometry of a standard 384 well (Figure 1b). The boundary condition of the sidewalls was set to no flux.The Partition Condition boundary condition (a built-in function in the CRE Module) defines different gas solubilities in neighboring materials (oil/media, etc.). Oxygen diffuses from the top surface [21% O_2_ saturated (i.e. oxygen solubility) of a given material] (Table 1) to the bottom surface. The boundary condition at the bottom surface (i.e. the cell layer) accounts for the O_2_ demand = -(*n* × OCR_Avg_ × *ρ*_cell_ × DF × POC)/(Km0 × S1 + POC), where OCR_Avg_ = 2.5 amol/cell/s is the average OCR of human body cells,^[58]^ *n* is the ratio between the OCR of a given cell type and OCR_Avg_, *ρ*_cell_ is cell density (3×10^4^ cells/well as the default density that leads to near confluence in a standard 384 well from cell seeding of the typical mammalian cells), DF is a dimensionless cell density factor ranging from 0 to 1, POC is the oxygen concentration at the cell layer defined by COMSOL at a time, Km0 is Michaelis-Menten constant of a given cell type (e.g. 5.6 mmHg for HUVEC), and S1 is the solubility of oxygen in cell (1.049 mM/atm).^[59]^ (i) Influence of diffusion barrier on establishing a supply-demand-balanced oxygen microenvironment (Figure 1b). In this section, we used *n* = 5, DF = 1 for the O_2_ demand. We set the diffusion barrier as PDMS elastomer, silicone oil (5 cSt), silicone oil (500 cSt) (Table 1) from left to right in Figure 1b, which gives hyperoxia, physioxia (i.e. about 5% O_2_ in homeostasis), hypoxia, and anoxia, respectively. (ii) Reconditioning of an established oxygen microenvironment against the change from supply or demand (Figure 1d,e,g). The geometry of the models in this section are the same as the ones shown in Figure 1b. We selected PDMS elastomer as the diffusion barrier material for the supply-dominated model (Figure 1d). For change from supply, we set the oxygen concentration at the top surface to: 0.2×O_2__inMedia (0.21 mol/m^3^) from 0 h to 8 h (Stage I), 0.8×O_2__inMedia from 8 h to 16 h (Stage II), and 0.7×O_2__inMedia from 16 to 24 h (Stage III). Here we used *n* = 3.75, DF = 1 for the O_2_ demand. For change from demand, we set the surface reaction at the bottom surface to: *n* = 4, DF = 1 from 0 h to 8 h (Stage I), *n* = 5, DF = 1 from 8 h to 16 h (Stage II), and *n* = 2, DF = 1 from 16 to 24 h (Stage III). Here we used O_2__inMedia for the O_2_ supply. We selected PS as the diffusion barrier material for the demand-dominated model (Figure 1e). For change from supply, we set the oxygen concentration at the top surface to: 1.0×O_2__inMedia from 0 h to 8 h (Stage I), 0.9×O_2__inMedia from 8 h to 16 h (Stage II), and 0.8×O_2__inMedia from 16 to 24 h (Stage III). Here we used *n* = 0.5, DF = 1 for the O_2_ demand. For change from demand, we set the surface reaction at the bottom surface to: *n* = 0.2, DF = 1 from 0 h to 8 h (Stage I), *n* = 0.4, DF = 1 from 8 h to 16 h (Stage II), and *n* = 0.8, DF = 1 from 16 to 24 h (Stage III). Here we used O_2__inMedia for the O_2_ supply. We selected silicone oil (5 cSt) as the diffusion barrier material for the supply-demand-balanced model (Figure 1g). For change from supply, we set the oxygen concentration at the top surface to: 1.0×O_2__inMedia from 0 h to 8 h (Stage I), 0.85×O_2__inMedia from 8 h to 16 h (Stage II), and 0.95×O_2__inMedia from 16 to 24 h (Stage III). Here we used *n* = 4, DF = 1 for the O_2_ demand. For change from demand, we set the surface reaction at the bottom surface to: *n* = 4, DF = 1 from 0 h to 8 h (Stage I), *n* = 6, DF = 1 from 8 h to 16 h (Stage II), and *n* = 2, DF = 1 from 16 to 24 h (Stage III). Here we used O_2__inMedia for the O_2_ supply. (iii) The kinetics of oxygen diffusion in multiliquid-phase microsystems (Figure 2). In these models, we set the O_2_ demand at the bottom surface the same with *n* = 14 and with DF = 1 (for 3×10^4^ cells/well) or DF = 1/3 (for 1×10^4^ cells/well) for different cell densities. We set the O_2_ supply at the top surface and the initial oxygen concentration in each layer based on the corresponding material (Table 1). The readout of oxygen level (O_2_, %) was extracted from the cell layer at the bottom surface over 48 h. The .txt data sheets were exported and plotted in Prism Graphpad. The 3D diffusion profiles were generated in COMSOL with a color-coded O_2_, % bar denoting 0% and 21% O_2_ in culture media. The figures were prepared in Adobe Illustrator.

### Measurement of POC and IOC

(i) Working principle of the optical oxygen sensor (for POC). The optical oxygen sensor system (Ohio Lumex) includes an oxygen meter [FireStingO_2_ fiber-optical oxygen meter (PS FSO2-2)], a temperature sensor [Teflon-coated and submersible, not shielded, Ø = 2.1 mm (PS TSUB21)], an oxygen mini sensor [Ø = 430 μm (OXB430)], and a computer installed with Firesting Logger software (Pyro Science). The measurement is based on quenching of near-infrared (NIR) fluorescence in the presence of molecular oxygen in the media. The quenching of fluorescence is described by Stern-Volmer relationship as *I*^0^/*I* = 1 + *K*_SV_[O_2_], where *I*^0^ and *I*, respectively, correspond to the fluorescence intensities in absence and presence of oxygen; *K*_SV_ is the Stern-Volmer constant, and [O_2_] is the concentration of oxygen in the sample. (ii) POC measurement with the optical oxygen sensor. The 384-well plate from cell culture was placed on a hot plate (40 °C, the set temperature) on the bench. An aluminum bar was applied between the well plate and the surface of the hot plate to enhance heat transfer. The temperature sensor was kept in a well (without cells) filled with 50 µl DI water and with the tip sitting on the bottom of the well to give accurate temperature compensation. The system was allowed to stabilize for at least 5 min before the calibration, measurement and data collection. Calibration of the optical oxygen sensor was done using the “2-point in water or humid air” mode with the steps as follows: #1) Prepare the two calibration liquids. Liquid A - air-saturated DI water for 100% O_2_ saturation ratio (or 21% O_2_); Liquid B - freshly prepared 1 wt% sodium sulfite (Na_2_SO_3_) (≥ 98%, Sigma Aldrich, S0505) aqueous solution for 0% O_2_ saturation ratio (or 0% O_2_). Give 5 min on Liquid B to reach the steady stage with 0% O_2_ before use. Add 50 µl of each liquid to a well on the well plate and prewarm; #2) Sterilize the oxygen sensor by submerging the sensor tip in 70% ethanol (Thermo Fisher Scientific, 64-17-5) in an eppendorf tube for 10 sec. Then thoroughly rinse the sensor tip with DI water and dry it in air using a rubber bulb; #3) Keep the oxygen sensor in Liquid A first until the oxygen readout curve is stabilized at 100% O_2_ saturation ratio (with minimal background fluctuation). Then switch to Liquid B to have the oxygen readout curve stabilized at 0% O_2_ saturation ratio. Thoroughly rinse the sensor tip with DI water and dry it in air using the rubber bulb; #4) Insert the oxygen sensor mounted on a linear translation stage (Siskiyou, MX130L) into a well through the oil overlay with the sensor tip resting on the cell layer at the bottom of the well to give an accurate readout of POC. Each recording of the POC lasts an equal length of time (e.g. 1 min) after the reading is stable. The replicate wells from the same cell type and condition are measured in a row without additional treatment of the sensor tip during switch. To switch to a different cell type or condition, the sensor tip is sterilized following Step #2 before the next measurement to avoid cross contamination; #5) After all measurements finished, clean the oxygen sensor following Step #2; #6) The recorded txt. data sheets (O_2_ % versus time) were pooled together for each condition in Excel and then plotted in Prism GraphPad. (iii) Working principle of the hypoxia dye (for IOC). The measuring of the hypoxia dye is based on the uptake of the dye molecules into live cells which fluoresce when experiencing a reduced oxygen level inside the cells (i.e. IOC) compared to 21% O_2_. Dead cells release the dye and lose fluorescence. A couple of common pitfalls from using this hypoxia dye need to be clarified: #1) The dye only provides a fluorescence readout that can be used to reflect/track the drop of IOC compared to 21% O_2_. The fluorescence intensity from a single condition on itself doesn’t tell any specific oxygen levels unless benchmarked to a direct oxygen level readout, e.g. POC measured at the cell layer (which is the protocol used in this work); #2) Moreover, the fluorescence intensity can vary with different operation parameters including the concentration of the dye in media and imaging conditions (e.g. the bottom material/thickness of the well plate, laser intensity, exposure time). A parallel comparison of the fluorescence intensity is valid only if the operation parameters are defined and maintained consistent; #3) At last, the brand name of the dye (Hypoxia Reagent) itself is apparently based on the conventional definition of “normoxia” with 21% O_2_, however, inaccurate and misleading.^[60]^ An increased fluorescence compared to the baseline with 21% O_2_ may indicate an oxygen level anywhere between hyperoxia and anoxia (Figure 1a,b). In other words, the hypoxia dye doesn’t only report hypoxia. (iv) IOC tracked by the mean fluorescence intensity of the hypoxia dye. Fluorescent imaging was performed on a Nikon Ti Eclipse inverted epifluorescence microscope (Nikon Instruments) with ×3 replicate wells from each condition to get the hypoxia dye signal at 4× magnification (to cover the cell layer from the whole well on a 384-well plate) with 485 nm/525 nm [Excitation (Ex)/Emission (Em)], maximum laser power, and 1 sec of exposure time. Wells with the same cell seeding but without hypoxia dye were imaged for control and background subtraction during image processing and analysis of the mean fluorescence intensity. The mean fluorescence intensity was extracted from an image (in 16-bit color depth, with and without hypoxia dye) using the “Analyze → Measure” function in Fiji ImageJ, with a range of interest (ROI) set to cover the cell layer but exclude the edges of a well. The mean fluorescence intensities of images with hypoxia dye were subtracted with the average of the mean fluorescence intensities from the control without hypoxia dye.

### Oxygen diffusion test of silicone oil

1 ml of culture media (DMEM + 10% FBS) was pipetted to a 5 ml centrifuge tube (polypropylene, Argos, T2076A). The oxygen sensor was submerged in the media with the tip being kept about 1 mm from the air/media interface (Figure S2). A stainless steel blunt needle (18G, SAI Infusion Technologies, B18-150) was connected to a N_2_ cylinder via silicone tubing (Tygon) and then kept close to the bottom of the centrifuge tube to perform N_2_ bubbling. The N_2_ flow rate was set at about 4 ml/min. N_2_ bubbling was let run for about 500 sec (∼8 min) to reach 0% O_2_ of the media. N_2_ bubbles got accreted at the air/media interface and were broken by keeping the needle in the bubble layer for a few seconds. After the N_2_ bubbles were removed the N_2_ gas trapped in the centrifuge tube was purged by a rubber bulb three times to create the air (i.e. no oil) condition or was directly replaced by silicone oil (3 ml) [Sigma Aldrich, 317667 (5 cSt), 378399 (1000 cSt)] added by pipette. The 3 ml of oil added on top of the media in the 5 ml centrifuge tube led to about 18 mm in the oil depth. Then the oxygen recovery was recorded with the optical oxygen sensor until the oxygen level reached about 10% O_2_. The recorded txt. data sheets (O_2_ % versus time) were plotted in Excel.

### UPLC-MS media analysis

The objective of this assay is to compare silicone oil and fluorinated oil treated media to control media for relative quantification of media composition change. The equipment is a tandem of a liquid chromatography (LC) system (Dionex UPLC) with Amino Acid columns (Kinetix Imtakt Intrada mixed mode Amino Acid, 3 x 150 mm) and a Mass Spectrometer [Thermo Q-Extactive (QE) Orbitrap]. Samples include control media (DMEM + 10% FBS + 1% Pen-Strep), silicone oil (5 cSt), and fluorinated oil (Fluorinert FC-40) (Sigma Aldrich, F9755) treated media (Figure 3a) for ×3 independent replicates of each condition. Samples were prepared with the steps as follows: #1) Add 300 µl precipitation mix (acetonitrile with formic acid and *d*6-GABA internal standard) to 9 Sirocco wells; #2) Add 100 µl each sample to each well and let sit for 2 min; #3) Push through the plate, dry under nitrogen, and resuspend dried samples in 100 µl 0.1% formic acid/water for amino acid runs; #4) Inject 2 µl of each sample in duplicate on QE using a top 5 ms/ms method with a scan range of 80-700 m/z. Also make 1 pooled quality control sample of equal molar concentration of all 9 samples (injected twice) for normalizing data during data analysis. Method run on UPLC Amino Acid - Solvent A (acetonitrile/0.3% formic acid); Solvent B (water 100 mM ammonium formate); Flow (0.5 ml/min); Column Temp (60 °C); Max Pressure Limit (7,000 psi); Running pressure (1,000 psi). Method run on QE - FULL MS / DD-MS. (TOPN) - Full MS: Microscans (1); Resolution (70,000); Automatic gain control (AGC) target (1e6); Maximum injection time (IT) (100 ms); Scan range (80 to 700 m/z); Spectrum data type (Profile). Microscans 1: Resolution (17,500); AGC target (1e5); Maximum IT (100 ms); Loop count (5); Isolation window (3.5 m/z); Normalized collision energy (NCE) (30.0); Spectrum data type (Centroid); Peptide match (off); Exclude isotopes (on); Dynamic exclusion (1.0 s). The following filters were used when analyzing the data using Compound Discoverer 3.0 (Thermo Scientific Metabolomics Software): #1) Base peak intensity of all samples > 1e5. This filter is for picking peaks that are above background noise level; #2) Masses for identified compounds had to be within 5 ppm of theoretical values; #3) ms/ms patterns of identified compounds had to match theoretical fragmentation patterns; #4) Percent coefficient of variations (CVs) for replicates within a given condition or control 6 replicates had to be < 20%; #5) Data was normalized using the x2 injection of the pooled quality control sample (Median Absolute Deviation). Meta data from Compound Discoverer 3.0 (filtered data only): #1) A total of 193 peaks (compounds) were aligned between the control and treated samples and passed all above filters; #2) Of the 193 compounds, 39 were identified using molecular weight and ms/ms fragment patterns using above filters; #3) For the silicone oil samples none of the identified compounds showed any difference in regulation in comparison with the control media samples. However, two of the unidentified compounds had adjusted p-values that were significant when compared to the control; #4) Similarly, for the Fluorinert FC-40 samples none of the identified compounds showed any difference in regulation in comparison with the control samples. Additionally, two unidentified compounds [different mass and retention time (RT) than for the silicone oil] had adjusted p-values that were significant when compared to the control; #5) An Excel report that has all the relevant information (relative intensity, compounds masses, names, statistics) was generated using the filtered data. #6) The HLB values of the identified molecules were obtained in Chemicalize (ChemAxon). The AUCs (i.e. area under the curve) of the top five lipophilic molecules with HLB ranging from 0 to about 6 were plotted in Prism GraphPad.

### Preparation of mono-culture plates

The mono-culture was established in (but not limited to) 384-well plates (Polystyrene, Tissue Culture Treated, Flat Bottom, 384-well plate, Corning 3701) with cell culture media corresponding to each cell type (Table S1), and with silicone oil or fluorinated oil (Fluorinert FC-40) overlay. The culture media and oil in this work were not deoxygenated prior to use. The plates were prepared in a sterile biosafety hood in air at room temperature (RT, ∼22 ^°^C). Specifically, to prepare a mono-culture plate include the following steps: #1) Add a certain volume (e.g. 15 or 35 µl/well) of fresh media (stored in a regular 4 ^°^C fridge, oxygen saturated) with and without the hypoxia dye (Invitrogen, Image-iT Green Hypoxia Reagent, Thermo Fisher Scientific, I14833) (1:1000 dilution in media); #2) Overlay the media with a certain volume [e.g. 0 (for no-oil control), 10 or 50 µl/well] of oil; #3) Prewarm the plate in a standard incubator [37 ^°^C, 18.6% O_2_, 5% carbon dioxide (CO_2_), 95% relative humidity (HR)] before cell seeding; #4) Prepare the cell stock at a specific concentration (e.g. 2000 or 6000 cells/µl) following the standard cell culture/passage protocol; #5) Pipette 5 µl of the cell stock to the media under oil to reach a target seeding density (e.g. 1×10^4^ or 3×10^4^ cells/well); #6) Keep the plate in a standard incubator up to 24 h or 48 h without media change. The plates were imaged and measured to get different readouts, e.g. cell viability, POC and IOC (see details in Measurement of POC and IOC above).

### Preprocessing of cells in mono-culture

(i) Preculture of *C. albicans. C. albicans* fungal cells (CMM16) were inoculated from a streaked plate in 2 ml yeast extract peptone dextrose (YPD) [1% yeast extract (BD Biosciences, 212730), 2% peptone (BD Biosciences, 211862), 2% dextrose glucose (Thermo Fisher Scientific, 215510)] glucose media and grown in an incubator at 30 ^°^C overnight. The fungal cells were measured on a spectrophotometer (Thermo Fisher Scientific, 335932) with optical density at a wavelength of 600 nm (OD_600_) and then converted to cell concentration. Serial dilution was performed to reach a final concentration of 6000 cells/μl in a culture media (Table S1) for further experiments. (ii) Isolation of neutrophils from whole blood. Primary human neutrophils were isolated from peripheral blood taken from healthy donors. All blood samples were drawn according to Institutional Review Boards (IRB)-approved protocols per Declaration of Helsinki at the University of Wisconsin–Madison. Peripheral neutrophils were isolated by negatively removing all contaminating cells using the MACSxpress Neutrophil Isolation Kit (Miltenyi Biotec, 130-104-434) and BD Pharm Lyse buffer (BD Biosciences, 555899) for red blood cell depletion, according to manufacturer’s instructions. Cells were washed and resuspended in appropriate media (Table S1) for further experiments.

### Preparation of mono-culture in PDMS microchannels

The PDMS microchannels (about 2500 μm in length, 600 μm in width, and 250 μm in height) (Figure S3) were prepared following a standard photolithography process and O_2_ plasma bound onto a chambered coverglass [Nunc Lab-Tek II Chambered Coverglass, #1.5 borosilicate coverglass (0.16-0.19mm), biocompatible acrylic adhesive, Thermo Fisher Scientific, 155382]. Collagen type I (Corning, 354249) solution prepared with 10× phosphate-buffered saline (PBS), cell culture media, and 0.5M NaOH at 3 mg/ml was added to the glass surface in the microchannels to create a thin layer of collagen coating prior to polymerization. After polymerization of the collagen coating at 37 °C, 30 μl of Caco-2 cells at a concentration of 15000 cells/μl in culture media (EMEM + 20% FBS) were seeded into each microchannel via passive pumping and cultured for 24 h. After 24 h cell culture the conditioned media (along with the suspension cells) was replaced with a fresh media containing hypoxia dye. The microchannels were then overlaid with a 2.5 mm depth of fluorinated oil (Fluorinert FC-40), followed by an additional 5 mm depth of silicone oil (1000 cSt). The chambered coverglass with the PDMS microchannels was kept in a standard incubator for 48 h without media change before the characterization and imaging on a microscope (see IOC tracked by the mean fluorescence intensity of the hypoxia dye).

### Reconditioning of the under-oil oxygen microenvironment

We performed this test by regulating mitochondrial respiration in UOMS cell culture. A 384-well plate of Caco-2 cells was prepared (see Preparation of mono-culture plates) by seeding 1×10^4^ cells/well in 20 µl of culture media (EMEM + 20% FBS) with 5 mm silicone oil (5 cSt) overlay. The cells were cultured for 12 h to reach a starting level of POC at about 15% O_2_. A mitochondrial stress assay kit (Agilent Technologies, Seahorse XFp, 103010-100) was used to manipulate the mitochondrial respiration and thus the OCR of the cells. Specifically, FCCP [carbonyl cyanide-4 (trifluoromethoxy) phenylhydrazone] collapses the proton gradient and disrupts the mitochondrial membrane potential. As a result, electron flow through the ETC (i.e. Electron Transport Chain) is uninhibited, and oxygen consumption by Complex IV reaches the maximum. A mixture of ROT (rotenone)/AA (antimycin A) inhibits Complex I and Complex III. This combination shuts down mitochondrial respiration. FCCP and the mixture of ROT/AA were reconstituted in DMEM culture media (not deoxygenated) to get a 50 µM and 25 µM compound solution, respectively. The POC was monitored in real time for 3 h by the optical oxygen sensor (see POC measurement with the optical oxygen sensor) with the culture plate set on a hot plate. The compound solutions were added to a well at a volume of 0.4 µl/dose [for a 1:50 v/v ratio (0.4 μl compound solution:20 μl media)] in a sequence (×4 doses of FCCP and ×2 doses of ROT/AA). To avoid introduction of air during the pipetting under oil, the pipet tip was prefilled with oil and then loaded with 0.4 µl of the compound solution. Compounds were added to the culture media with the prefilled oil kept in the tip.

### Colorimetric analysis of pH in cell culture with and without oil overlay

The test was performed with mono-culture of Caco-2 cells (see *Preparation of mono-culture plates*) compared to a no-cell control (Figure S13). The culture media (EMEM + 20% FBS) was supplemented with a pH indicator (phenol red, 30 μM). A pH color chart was obtained using a pH meter (Model 225, Electrode Model 300731.1, Denver Instrument) and hydrochloric acid (1 M) titration. 10 ml of the media was added to a 50 ml centrifuge tube (Falcon, 734-0451). Hydrochloridric acid was added to the media to reach a specific pH reading. 20 μl of the pH-adjusted media was pipetted to a well of a 384-well plate and then pictured (iPhone 6s) against a white background. For the culture and control plates, each condition has at least ×3 replicates. The plates were pictured before and 24 h after incubation in a standard incubator with different exposure times (i.e. 0 min, 30 min, and 100 min) in air at room temperature. The color of the media in comparison with the pH color chart indicates the pH level of the media in a tested condition, reflecting a reduced CO_2_ diffusion from media to ambient (∼0.04% CO_2_) with the oil overlay.

### Live-dead staining and cell viability analysis

Cells were stained with 2 μM calcein AM (Thermo Fisher Scientific, L3224), 1 μM propidium iodide (Thermo Fisher Scientific, P1304MP) and Hoechst 33342 (Thermo Fisher Scientific, H3570) in appropriate culture media (Table S1), following 24 h of culture to obtain live, dead and nuclei counts, respectively. Staining solution was prepared at 2× immediately before staining and diluted in culture media within wells to make 1× solution. Cells were incubated with staining solution for 20 min in a standard incubator prior to imaging. Fluorescence images were obtained on a Nikon Ti microscope and kept at 37 ^°^C and 5% CO_2_ via an on-stage incubator (Bold Line, Okolab) during imaging. A minimum of ×3 replicate wells with live cells were averaged for viability analysis. The dead-channel images were threshold processed using the “Image → Adjust → Threshold (Default)” function in Fiji ImageJ. Threshold adjustment was carried out on the entire image including the edges of the well. Minimal adjustment might be applied manually if necessary to get the optimal threshold-processed images that loyally distinguish and pick the cells. With the seeding densities in this study (i.e. 1×10^4^ or 3×10^4^ cells/well of a 384-well plate), cell clumping could happen. The cells in a clump can not be effectively identified by the software as separate entities, which causes negative errors on cell count. The binary watershed function “Process → Binary → Watershed” in Fiji ImageJ was chosen to process the image further so that the clumps were broken into smaller particles to give a more accurate cell count of the true amount of cells by the software. On the processed images, particle analysis was carried out using the “Analyze → Analyze Particles…” function in Fiji ImageJ, which gives the cell count (of the dead cells in this case) in the output. In theory, the total cell count (including both live and dead cells) can be obtained by analyzing the nuclei-channel images. However, cell clumping became a dominant effect on the nuclei-channel images, which led to a severely underestimated number of the total cell count. Provided such a limit, we used the total number of cells in each seeding density (i.e. 1×10^4^ or 3×10^4^ cells/well) instead to calculate the cell viability, following the equation: cell viability = live cells/total cells × 100% = (total cells - dead cells)/total cells × 100%.

### Preparation of co-culture plates

The co-culture was performed in 384-well plates [Glass Bottom (#1.5, 0.170 ± 0.005 mm, high performance coverglass), Black polystyrene frame, Tissue Culture Treated, Flat Bottom, 384-well plate, Cellvis, P384-1.5H-N] with the steps as follows: (i) Isolation of intestinal crypts and generation of colon organoids. All human studies were approved by an IRB at University of Wisconsin-Madison (Protocol# 2016-0934). Intestinal crypts were isolated from colon removed for nonmalignant etiology using a previously established protocol.^[61]^ Isolated crypts were embedded in Matrigel (Basement membrane, Corning, 354248), and cultured in 24-well plates (Polystyrene, Nunc, Non-Treated Multidishes, Thermo Fisher Scientific, 144530) in intestinal stem cell media (Table S1). Colon organoids were split 1:2 to 1:4 every 2 to 3 days using a previously established protocol.^[62]^ (ii) Culture of colon monolayer. Based on the established protocol,^[62]^ intestinal stem cell media was removed from the 24-well plates containing colon organoid cultures, 500 μl of 0.5 μM ethylenediaminetetraacetic acid (EDTA) in PBS was added to each well and Matrigel plugs containing the organoids were harvested. After centrifugation at room temperature for 3 min at 300×g, supernatant was removed. 1 ml of 2.5 mg/ml trypsin (Sigma Aldrich, T4549) was added and incubated for 2 min in a 37 ^°^C water bath. Trypsin was neutralized using media [Advanced DMEM/F12, Thermo Fisher Scientific, 12634-010] + 20% FBS. The organoids were broken down into small fragments by pipetting vigorously using filtered 1000 μl pipette 60 times. 20,000 organoid fragments (500 fragments/μl) were seeded onto a 384-well plate coated with Matrigel that was diluted 1:40 in intestinal stem cell media. Media was changed every 2 to 3 days (stem cell media). After 7 days, a confluent monolayer was formed and the media was switched to 40 μl/well of differentiation media (for 4 mm in media depth) (Table S1) and 50 μl/well of oil (silicone oil, 5 cSt) (for 5 mm in oil depth) was overlaid. Bacterial co-cultures were performed on the following day. (iii) Preculture and fluorescent tagging of gut anaerobic bacteria. *B. uniformis* (DSM 6597) was cultured in an anaerobic chamber (Coy Labs) with 83% N_2_, 2% hydrogen (H_2_) and 15% CO_2_ at 37 °C in Anaerobe Basal Broth (ABB, Oxoid) for all steps except conjugation. During the conjugation procedure, *B. uniformis* was cultured in Brain Heart Infusion Broth (BHIS, Sigma Aldrich). The conjugation donor *E. coli* strain BW29427 was obtained from the *E. coli* Genetic Stock Center (CGSC). This strain was grown aerobically in Luria Bertoni (LB, Sigma Aldrich) media containing 25 mM of 2,6-Diaminopimelic acid (DAP, Sigma Aldrich). The *E. coli* strain BW29427 was transformed with the plasmid pWW3515 that harbored the fluorescent reporter mCherry expressed from the BfP1E6 promoter.^[63]^ The pWW3515 plasmid encodes the IntN2 tyrosine integrase, which mediates recombination between the *attN* site on the plasmid and one of the two *attBT* sites on the *B. uniformis* chromosome.^[64–67]^ Following the transformation, single colonies were inoculated into LB containing 100 µg/ml carbenicillin (Carb, IBI Scientific) and DAP and incubated at 30 °C overnight. Cell pellets were collected by centrifugation at 4,000 rpm for 5 min and washed with fresh LB media. The *E. coli* cells were then combined with the *B. uniformis* culture (OD_600_ = 0.5∼0.6) at a donor:recipient ratio (v/v) of 1:10. The cell mixture was pelleted, resuspended in 0.2 ml BHIS and then spotted on BHISAD (BHIS + 10% ABB + DAP) agar plates and incubated anaerobically at 37 °C for 24 h. The cell lawns were scraped and resuspended in BHIS and plated as serial dilutions on BHISAGE plates [BHIS supplemented with 10% ABB, 200 µg/ml gentamicin (Sigma Aldrich) and 25 µg/ml erythromycin (Sigma Aldrich)] and incubated anaerobically at 37 °C for 2 days. The engineered strain was verified by colony PCR. Bacterial inoculum onto the 384-well plate was captured in exponential phase, diluted to an OD_600_ of 0.1 in the culture media (e.g. the differentiation media for primary colon epithelium). (iv) Co-culture of colon monolayer with gut anaerobic bacteria. The bacterial inoculum stock (OD_600_ = 0.1) was prepared in the anaerobic chamber and transported in an anaerobic Hungate tube (VWR, 100484-346). The following inoculation was performed in air in a sterile biosafety hood. An eppendorf tube (Polystyrene, 0.6 ml) was prefilled with 300 μl of silicone oil (5 cSt). 100 μl of the bacterial inoculum was transferred from a Hungate tube to under oil in the eppendorf tube using a syringe (1 ml, Luer-lok Tip, BD Biosciences, REF 309628). Bacteria were added to the media above the colon monolayer by pipette at 1:20 v/v ratio (2 μl bacteria:40 μl media) carefully so as not to disrupt the oil overlay. Mono-culture conditions (including bacteria only and colon monolayer only) with and without oil overlay were prepared in parallel as control. Following the inoculation, the culture plates were kept in a standard incubator and cultured for 24 h.

### Downstream characterizations in co-culture

(i) RT-qPCR (colon monolayer). After 24 h of co-culture, the overlaying oil and media were carefully removed and each well was washed with Dulbecco’s PBS (Thermo Fisher Scientific, 14190250). To each well, 20 μl of Buffer RLT plus + β-mercaptoethanol (Sigma Aldrich, M3148) was added to lyse the cells. RNA was isolated using Qiagen RNeasy Plus Micro Kit (Qiagen, 74034) according to the manufacturer’s protocol and quantified with a pico chip on an Agilent Bioanalyzer (Santa Clara, CA). Reverse transcription was conducted with the RNA to cDNA kit (4387406, ThermoFisher Scientific). A preamplification step that amplifies the amount of cDNA available for downstream RT-qPCR analysis was conducted using SsoAdvanced PreAmp Kit (BioRad, 1725160). RT-qPCR was performed using TaqMan Gene expression assays (Thermo Fisher Scientific) with LightCycler 480 Probes Master (Roche Diagnostics). A 20 μl total reaction volume was used with 1 μl of 20× TaqMan primer-probe mix, 10 μl of 2× LightCycler 480 Probes Master mix, and 9 μl of cDNA diluted in RNase-free water. RT-qPCR amplification was monitored using a LightCycler 480 (Roche Diagnostics). qPCR was performed on 6 target genes and 3 reference genes (Table S2). The reference genes were selected based on constitutive and stable expression across sample types. After incubation at 95 °C for 10 min, the reactions underwent 45 cycles as follows: 10 sec at 95 °C, 30 sec at 60 °C, and 1 sec at 72 °C. Genes with Ct ≥ 35 were excluded. The Ct values were normalized to the reference genes (GAPDH, HPRT and RPLP0). Quantification of results (Figure 6f) are presented as: ΔCt = [CtGene-mean(CtGAPDH, CtHPRT, CtRPLP0)] so that positive values represent low expression and negative values represent high expression compared to the reference genes. (ii) Immunostaining (colon monolayer). The cells on a 384-well plate were fixed first by adding 40 μl of 4% paraformaldehyde (PFA) to each well and then incubated at room temperature for 10 min. After the PFA solution was removed by pipette, the fixed cell layer was washed with 100 μl of 1× PBS for three times. After fixation, the cells were permeabilized by adding 40 μl of 0.1% Triton X-100 to each well and then incubated at room temperature for 5 min. After the Triton solution was removed by pipette, the permeabilized cell layer was washed with 100 μl of 1× PBS for three times. After permeabilization, a blocking buffer consisting of 3% bovine serum albumin (BSA), 5% human serum, and 0.1 M glycine in 0.1% PBST (1× PBS with 0.1% added Tween 20) was added to each well at a volume of 80 μl. The plate was stored in a cold room overnight at 5 ^°^C to prohibit nonspecific binding of antibodies in the following staining process. Before staining, the blocking buffer was removed by pipette and the wells were washed with 100 μl of 1× PBS three times. A staining buffer was prepared utilizing 3% BSA in 0.1% PBST and 10% of a 1% Tween 20 solution at a 1:25 or 1:50 concentration with a primary antibody [(DAPI, 1:2500 dilution, Thermo Fisher Scientific, D3571), (F-actin, 1:500 dilution, Thermo Fisher Scientific, T7471), (Rabbit polyclonal anti-FABP1, 1:25 dilution, Sigma Aldrich, HPA028275), (Mouse monoclonal anti-MUC2, 1:50 dilution, Santa Cruz Biotechnology, sc-515032), (Villin, Novus Biologicals, NBP2-53201), (Rabbit polyclonal anti-ZO-1, 1:25 dilution, Thermo Fisher Scientific, 61-7300), and (Mouse monoclonal anti-E-cadherin, 1:25 dilution, BD Biosciences, 610182)]. The staining buffer and accompanying antibody were added at a volume that was just large enough to cover the cell layer at the bottom of each well (∼10 μl). The plate was stored in a cold room overnight at 5 ^°^C. Removal of the buffer was followed by two washes of 0.1% PBST of about 100 μl/well. A secondary antibody [488-anti-mouse, 1:250 dilution, Abcam, ab150105) or (647-anti-rabbit, 1:250 dilution, Abcam, ab150075)] depending on the compatibility of the original primary antibody was mixed with the staining buffer at 1:250 concentrations along with a 1:5000 concentration (sometimes 1:3000 depending on available product) of DAPI to stain the nuclei and a 1:500 concentration of rhodamine phalloidin to stain the (F-actin) cytoskeleton. This final staining solution was added to each well at a volume of 10 μl/well and then the plate was incubated for 1 h at room temperature while protected from light using aluminium foil. Following the incubation, the wells were washed with 0.1% PBST for two times before imaging. (iii) Fluorescent/confocal microscopy. The fluorescent images and videos were taken on a Nikon Ti Eclipse at 4×, 15× (10× objective with the 1.5× tube lens), and 30× (20× objective with 1.5× tube lens) magnifications with bright field and fluorescent channels, including 390 nm/440 nm [Excitation (Ex)/Emission (Em)] for nuclei (DAPI); 485 nm/525 nm for hypoxia dye (Image-iT Green Hypoxia Reagent), live dye (calcein AM), goblet cells (MUC2), microvilli (Villin); 560 nm/607 nm (Ex/Em) for dead dye (propidium iodide), mCherry, and cytoskeleton (F-actin); and 648 nm/684 nm for tight junction (ZO-1). Maximum laser power was applied if not stated otherwise. 3D confocal images were acquired with a Nikon A1-Si laser-scanning confocal microscope (Nikon Instruments). (iv) Bacteria count. The 30× magnification fluorescent images were threshold processed using the “Image → Adjust → Threshold (Triangle)” function in Fiji ImageJ. And then the bacterial cells were picked up and counted using the “Process → Find Maxima…” function in Fiji ImageJ. A prominence threshold was applied to each image to reach an optimal pickup of the singal (i.e. cells) against the noise (i.e. background). The bacteria count results were averaged with a minimum of ×3 replicates of each condition and plotted in Prism GraphPad. Note that *B. uniformis* stably express the mCherry proteins at low density (e.g. at early time points after inoculation). However, at 24 h, *B. uniformis* grew into a highly dense layer in co-culture, mostly losing their fluorescence due to the repressed expression of mCherry in low oxygen levels. The bacteria showed up in the bright-field as bright dots (i.e. the pole of the rod-shaped bacteria) with a clear contrast against the background, squirming around in Brownian motion. The bright-field images were threshold processed and rendered with pseudo red color to visualize the bacteria (Figure 6e), and used instead in the bacteria count for *B. uniformis* at 24 h in co-culture.

### Cell line authentication

(i) The mammalian cell lines (Caco-2, MDA-MB-231, HUVEC, and THP-1) were authenticated using short tandem repeat (STR) analysis. The STR analysis was performed with a cell pellet of about 2 million cells spun down with the media removed. The results were compared to an online database (e.g. Lonza, ATCC) to confirm the identity of a cell line. (ii) Sanger sequencing (Functional Biosciences, funding NIH) of 16s rRNA gene from colony picks of a single morphology quadrant streak was used to confirm identity of the bacteria (*B. uniformis*) in the co-culture experiment. 27 Forward (2_27F universal 16s rRNA gene forward primer) and 1492 Reverse (2_1492R universal 16s rRNA gene reverse primer) were used. Blast returns *B. uniformis* as top match in both cases, with 99.58% and 100% identity.

## Acknowledgements

C.L. conceived the AROM (i.e. autonomously regulated oxygen microenvironments) method and designed the research. C.L., G.M.W., and M.H. performed COMSOL Multiphysics simulation of oxygen diffusion in multiliquid-phase microsystems. C.L. designed and performed the UPLC-MS media analysis and data visualization with assistance from M.H., M.C.P.H., and C.O.S.. C.L. designed and performed the POC and IOC measurements and data analysis, visualization with assistance from M.H., J.L., Y.F., H.H., and J.S.. C.L. designed and performed the pH analysis in UOMS cell culture. C.L., M.H., K.P., B.C., and J.F. performed the cell culture/co-culture experiments with assistance from J.L., Y.F., H.H., and J.S.. J.F. constructed the fluorescently tagged *B. uniformis* strain, and B.C. carried out the pre-culture of bacteria with R.L.C. vetting the initial bacterial culture. C.L., M.H., and K.P. performed cell viability, IFS and qPCR characterizations. C.L., M.H., and B.C. prepared the cell line authentication. D.J.B., O.S.V., and C.L. supervised experimental design, data analysis, and data presentation. C.L., D.J.B., M.H., and O.S.V. wrote the manuscript and all authors revised it. This work was supported by NSF EFRI-1136903-EFRI-MKS, NIH R01 CA247479, NIH R01 AI154940, NIH R01 EB010039, NIH R01 CA185251, NIH R01 CA186134, NIH R01 CA181648, NIH P30CA014520, EPA H-MAP 83573701, American Cancer Society IRG-15-213-51 (Beebe Lab), and NIH R35GM124774 (Venturelli Lab). We thank Dr. Christopher Hartleb at the University of Wisconsin (UW)-Steven Points for demonstrating the (FireStingO_2_) optical oxygen sensor system, Dr. Evie Carchman from the Department of Surgery and UWCCC Biobank at the UW-Madison for the supply of colon tissue, McClean Lab from the Department of Biomedical Engineering at the UW-Madison for the supply of *C. albicans* (CMM16) sample, Huttenlocher Lab from the Department of Medical Microbiology and Immunology at the UW-Madison for the performance of blood draw. Small molecule relative quantitation by LC/MS/MS was done at the UW School of Pharmacy, Analytical Instrumentation Center (AIC) Mass Spectrometry Facility. We thank Molly C. Pellitteri Hahn and Cameron O. Scarlett in the AIC for their valuable contributions. We also thank the UW Translational Research Initiatives in Pathology laboratory (TRIP), supported by the UW Department of Pathology and Laboratory Medicine, UWCCC (P30 CA014520) and the Office of The Director-NIH (S10OD023526) for use of its facilities and STR Analysis services to perform cell line authentication, and the UW Optical Imaging Core for the support on the 3D confocal imaging. At last, we thank Dr. Jose M. Ayuso (Beebe Lab) for the introduction and early discussion upon the hypoxia dye in cell culture, and Mr. Duane S. Juang (Beebe Lab) for the inspiring conversation on the oxygen carrier function of fluorinated oil.

