## Supplementary material for "Under-Oil Autonomously Regulated Oxygen Microenvironments: A Goldilocks Principle-Based Approach For Microscale Cell Culture": N/A

Dr. C. Li, Prof. D. J. Beebe
Carbone Cancer Center
University of Wisconsin-Madison
Madison, WI 53705, USA

M. Humayun, J. Li, Y. Feng, H. Hefti, Prof. D. J. Beebe
Department of Biomedical Engineering
University of Wisconsin-Madison
Madison, WI 53705, USA

Prof. G. M. Walker
Department of Biomedical Engineering
University of Mississippi
University, MS 38677, USA

Dr. K. Y. Park
Department of Surgery
University of California San Francisco
San Francisco, CA 94143, USA

B. Connors, Dr. J. Feng, Dr. R. L. Clark, Prof. O. S. Venturelli
Department of Biochemistry
University of Wisconsin-Madison
Madison, WI 53706, USA

B. Connors, Prof. O. S. Venturelli
Department of Chemical and Biological Engineering
University of Wisconsin-Madison
Madison, WI 53706, USA

M. C. Pellitteri Hahn, Dr. C. O. Scarlett
Analytical Instrumentation Center-Mass Spec Facility
School of Pharmacy
University of Wisconsin-Madison
Madison, WI 53705, USA

J. Schrope
School of Medicine and Public Health
University of Wisconsin-Madison
Madison, WI 53726, USA

Prof. O. S. Venturelli
Department of Bacteriology

University of Wisconsin-Madison
Madison, WI 53706, USA

Prof. D. J. Beebe
Department of Pathology and Laboratory Medicine
University of Wisconsin-Madison
Madison, WI 53705, USA


**Table S1.** Compiled information of cell types and culture media.

| **Cell type (tissue origin)** | **Name** | **Culture media** |
|---|---|---|
| Endothelium (blood vessel) | HUVEC (human umbilical vein endothelial cell) | Endothelial basal medium-2 (EBM-2) (Lonza, 0019086) + 10% fetal bovine serum (FBS) (Thermo Fisher Scientific, 10437010) + 1% Penicillin-Streptomycin (Pen-Strep) (Thermo Fisher Scientific, 15070063) |
| Epithelium (colon cancer) | Caco-2 (cancer coli-2) | Eagle's minimal essential medium (EMEM) (Sigma Aldrich, M4655) + 20% FBS + 1% Pen-Strep |
| Epithelium (breast cancer) | MDA-MB-231 | Dulbecco's Modified Eagle's medium (DMEM) (Thermo Fisher Scientific, 11960051) + 10% FBS + 1% Pen-Strep |
| Fibroblast (normal) | Colon fibroblasts | Fibroblast media (ScienCell, C2301), 384-well plates coated with gelatin solution (Thermo Fisher Scientific, S25335) at 37 °C for 15 min and then aspirated. |
| Fibroblast (tumor-associated) | CAF (cancer-associated fibroblasts) (breast) | DMEM + 10% FBS + 1% Pen-Strep |
| Blood cells (monocytes) | THP-1 | Roswell Park Memorial Institute (RPMI) 1640 (Thermo Fisher Scientific, 11875085I) + 10% FBS + 1% Pen-Strep + 1% lactose |
| Blood cells (neutrophils) | Isolated from whole blood | EBM-2 + 5% FBS + 1% Pen-Strep |
| Primary epithelium (colon) | Colon organoids | Intestinal stem cell media [45% L-WRN conditioned media,[1] 45% midgut media,[2] 10% FBS, 50 ng/ml epidermal growth factor (EGF), 500 nM A-83-01, 10 µM SB202190, 10 nM $[Leu^{15}]$-Gastrin-1, 1 mM N-Acetylcysteine, 10 µM Y-27632, 2.5 µM CHIR99021, 2.5 µM Thiazovivin, and 100 µg/ml Primocin] |
| | Colon monolayer | Intestinal stem cell media (see above)<br><br>Differentiation media (5% L-WRN conditioned media, 85% midgut media, 10% FBS, 50 ng/ml EGF, 500 nM A-83-01, 10 nM $[Leu^{15}]$-Gastrin-1, 1 mM N-Acetylcysteine) |
| Fungi | *Candida albicans* (*C. albicans*, CMM 16 PES1 mutant) | RPMI 1640 |
| Bacteria | mCherry-labelled *Bacteroides uniformis* (*B. uniformis*, DMS 6597) | Anaerobe Basal Broth (Oxoid, CM0957)<br>Brain Heart Infusion Broth (Sigma Aldrich, 53286) (for conjugation) |

**Table S2.** The panel of genes in RT-qPCR and related protein function.

| Gene | | Protein function | Source |
|---|---|---|---|
| Proliferation | MKI67 | Cell proliferation marker | Thermo Fisher Scientific, Hs04260396_g1 |
| Differentiation | Axis inhibition protein 2 (Axin2) | A surrogate marker of intestinal stem cell activity (targeting Wnt signaling pathway) | Thermo Fisher Scientific, Hs00610344_m1 |
| | Trefoil factor 1 (TFF1) | An enterocyte marker (stabilization of mucus layer, healing of the epithelium) | Thermo Fisher Scientific, Hs00907239_m1 |
| | Sucrase-isomaltase (SI) | An enterocyte marker (digestion of dietary carbohydrates) | Thermo Fisher Scientific, Hs00356112_m1 |
| | Villin | Microvilli marker | Thermo Fisher Scientific, Hs01031739_m1 |
| | Mucin 2 (MUC2) | Goblet cell marker (epithelial lining) | Thermo Fisher Scientific, Hs03005103_g1 |
| Housekeeping (Reference genes) | GAPDH | N/A | Thermo Fisher Scientific, Hs01922876_m1 |
| | HPRT | N/A | Thermo Fisher Scientific, Hs02800695_m1 |
| | RPLP0 | N/A | Thermo Fisher Scientific, Hs99999902_m1 |

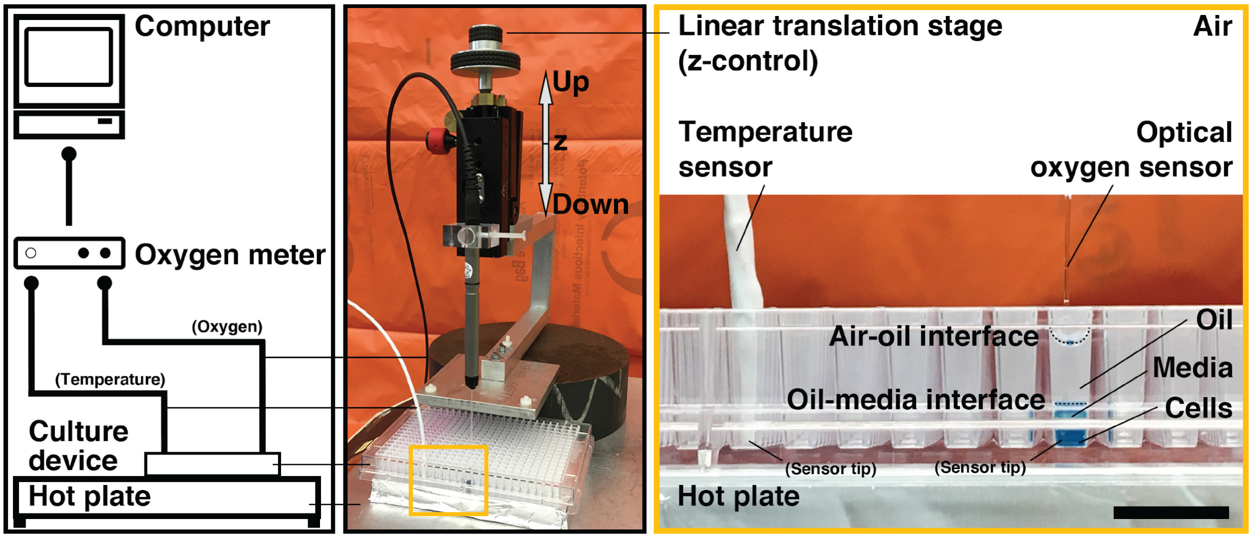


**Figure S1.** Camera pictures show the setup of UOMS in air with a 384-well plate and the optical oxygen sensor system for the measurement of POC. The well contains 20 µl of media (with blue food color for visualization), overlaid with 50 µl of silicone oil (5 cSt). For a standard 384 well, 10 µl leads to about 1 mm in depth. In UOMS cell culture, cells are seeded on the bottom of a well. The oxygen sensor is mounted on a linear translation stage for accurate position control in z-direction, resting on the cell layer during the measurement. The temperature sensor is submerged in a spare well filled with deionized water (50 µl) for real-time temperature compensation. Scale bar, 1 cm.

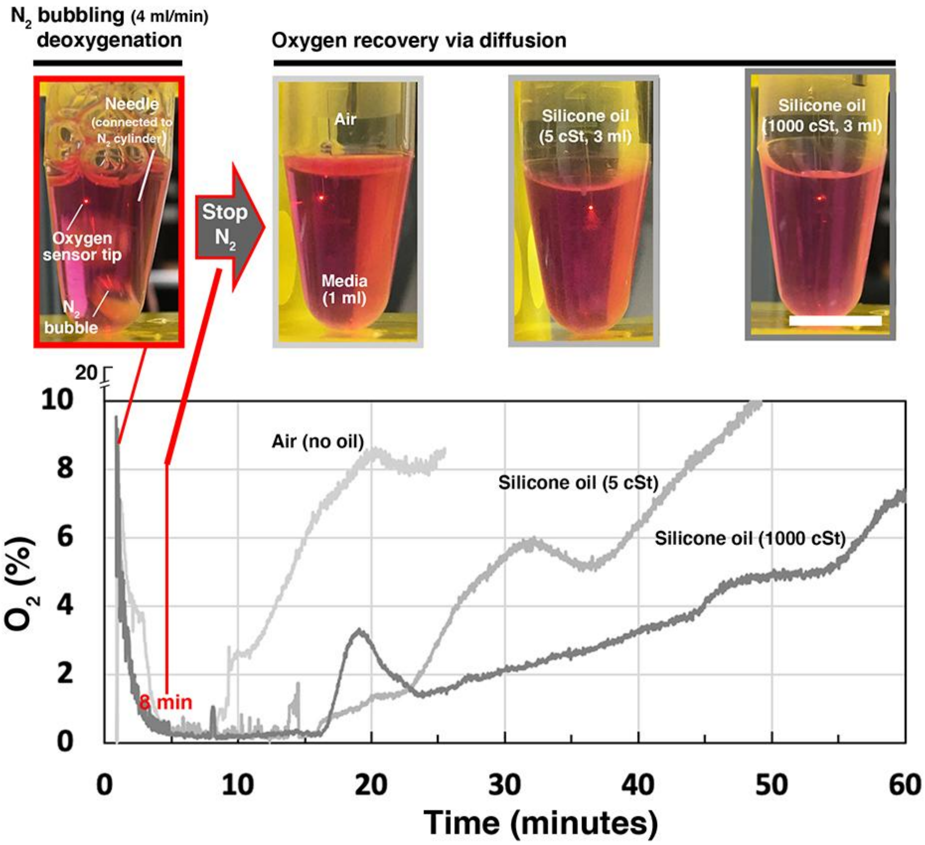


**Figure S2.** Oxygen diffusion test of silicone oil with different viscosities. Cell culture media (DMEM + 10% FBS) was deoxygenated to 0% $O_2$ by $N_2$ bubbling at a gas flow rate of about 4 ml/min. $N_2$ bubbling stopped at around 8 min. The media was overlaid with no oil (purged with air) or silicone oil in 5 cSt and 1000 cSt, respectively. The oxygen recovery process was recorded with the oxygen sensor tip kept at about 1 mm below the air/media (or oil/media) interface until it reached about 10% $O_2$. 3 ml of oil added on top of the media in the 5 ml centrifuge tube led to about 18 mm in the oil depth. Note that the $O_2$ signal fluctuations recorded in the conditions with oil overlay were caused by the metastability and spontaneous adjustment of the oil/media meniscus during measurements. Scale bar, 10 mm.

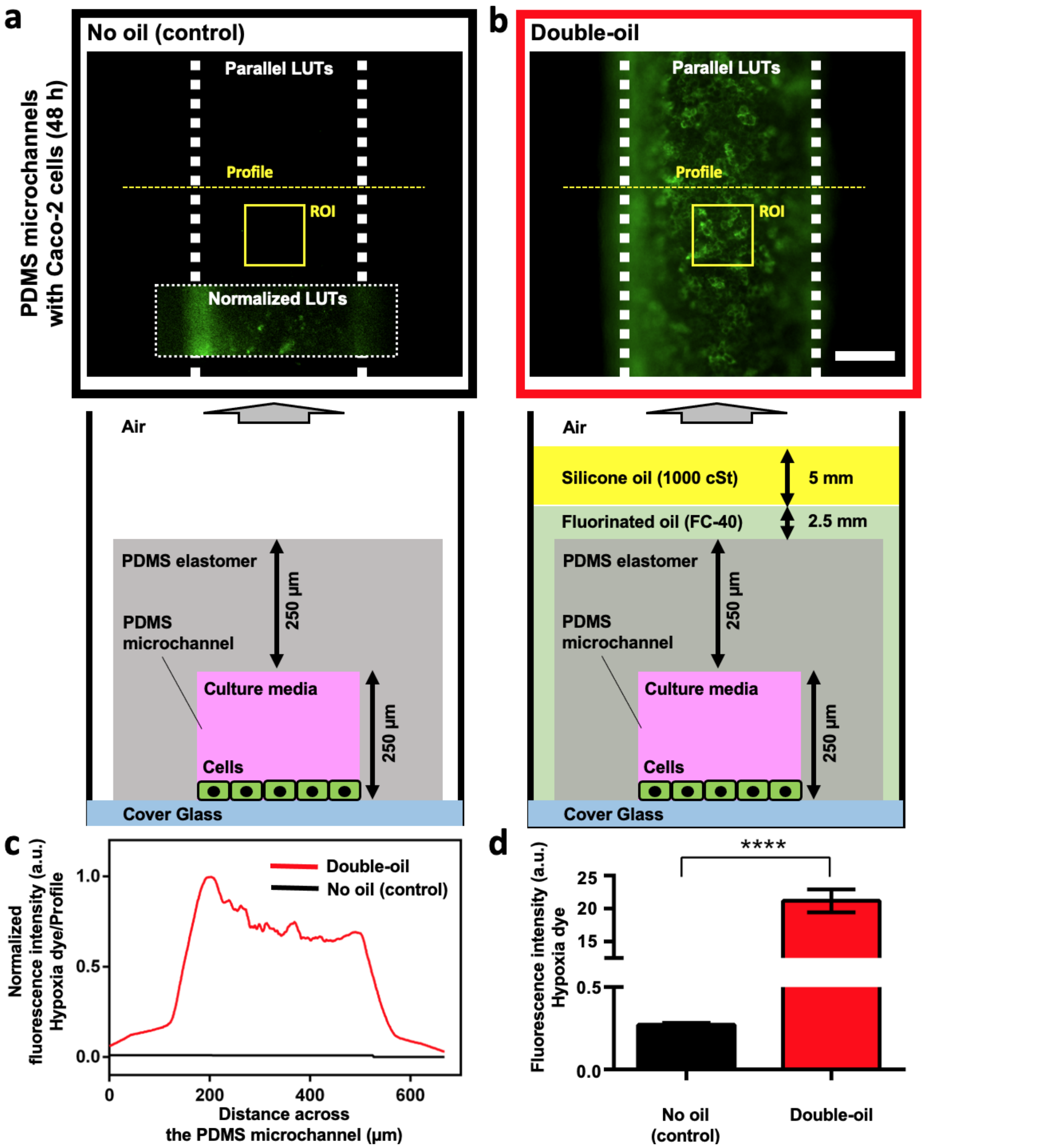


**Figure S3.** Comparison of hypoxia generation in PDMS microchannels with and without oil overlay. a) and b) The fluorescent images of hypoxia dye from each microchannel [no oil (control), left; double-oil, i.e. fluorinated oil (Fluorinert FC-40) + silicone oil (1000 cSt), right] with a confluent (Caco-2) cell monolayer (cultured for 48 h). Parallel LUTs (with exposure time of 500 ms), were applied for the comparison of fluorescence intensity. [Inset, no oil (control)] A fluorescent image with normalized LUTs to visualize the cells in the microchannel. The white dashed lines indicate the boundary of the microchannels. The channel dimensions are about 2500 µm in length (not fully shown in the images), 600 µm in width, and 250 µm in height. Scale bar, 200 µm. The schematic under each microchannel shows the cross section of the microchannels perpendicular to the length direction. c) The profiles of fluorescence intensity (normalized) across the microchannels (the yellow dashed lines in (a). d) The bar graph of IOC (fluorescence intensity of hypoxia dye) of each microchannel. The ROIs are shown by the yellow boxes in (a). Error bars, mean ± s.d. ****$P \leq 0.0001$.

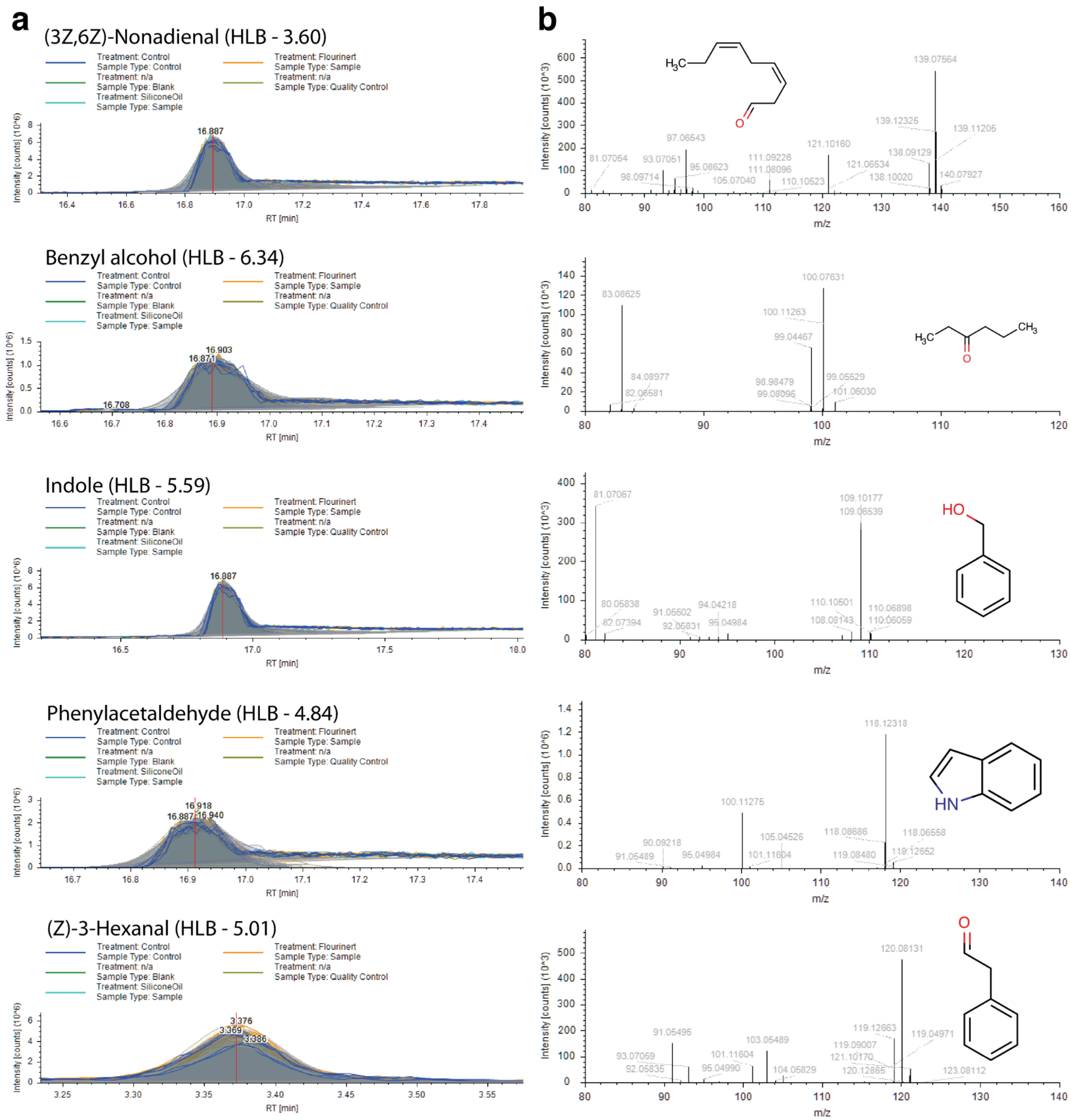


**Figure S4.** Raw UPLC-MS media analysis results. a) EICs (raw data, unsmoothed) of the five identified lipophilic compounds with their HLB values. Intensity of the signal at a chosen mass-to-charge (m/z) value is plotted against the retention time (RT). "Treatment: Control" is for the internal standard condition. See Experimental Section for details of the other conditions. b) ms/ms fragments spectra of each compound.

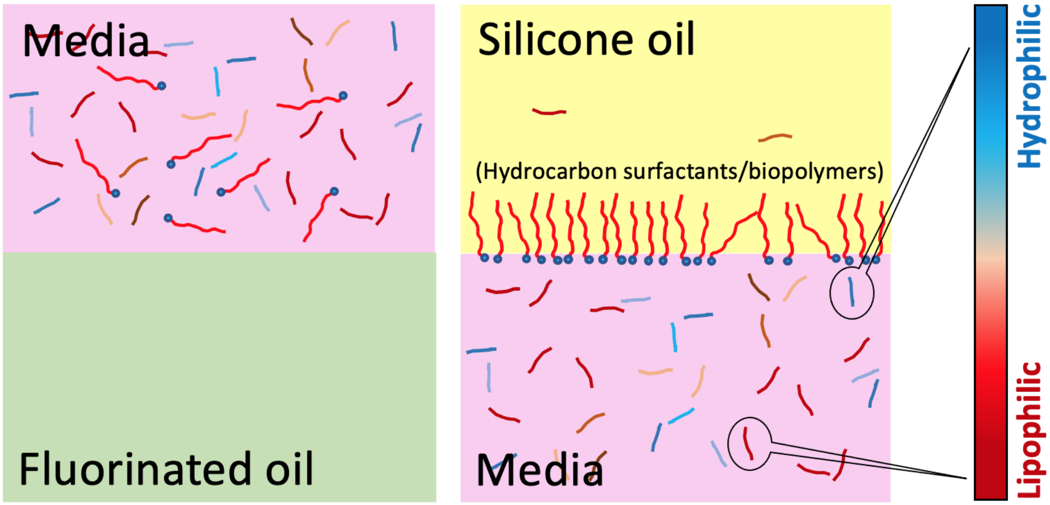


**Figure S5.** The proposed mechanisms for high retention of lipophilic molecules in media-fluorinated oil and media-silicone oil conditions.

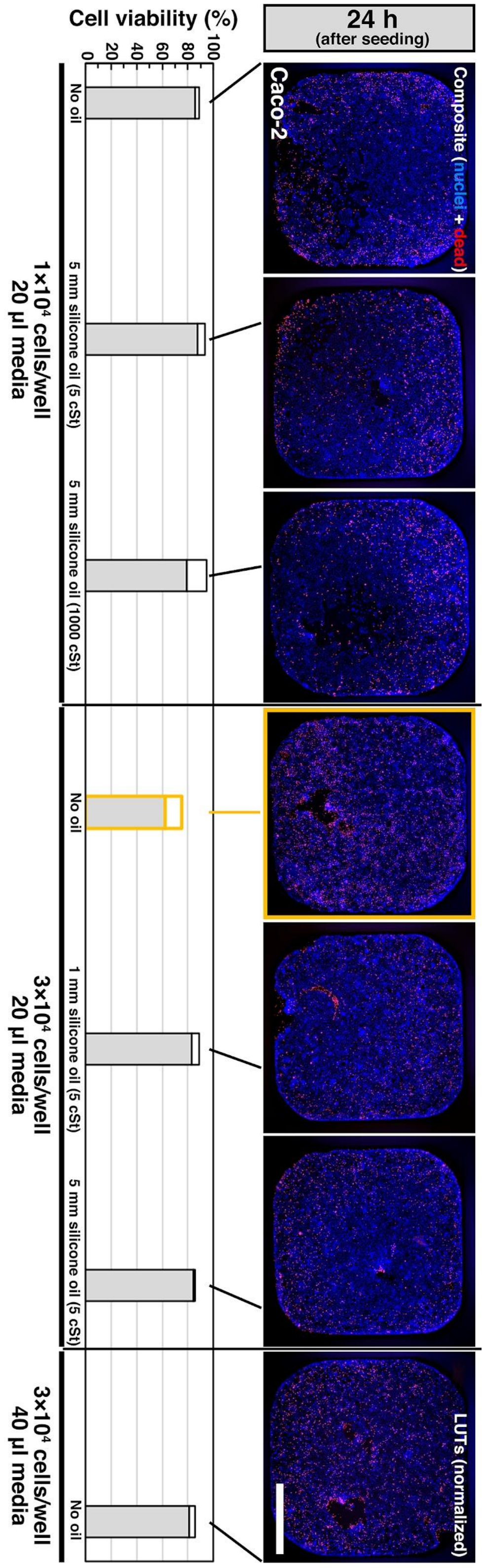


**Figure S6.** Cell viability of Caco-2 cultured with and without oil (silicone oil, 5 cSt) overlay (24 h after cell seeding with hypoxia dye). For a standard 384 well, 10 µl leads to about 1 mm in depth. The fluorescent images were all processed with normalized LUTs for visualization. Scale bar, 1 mm.

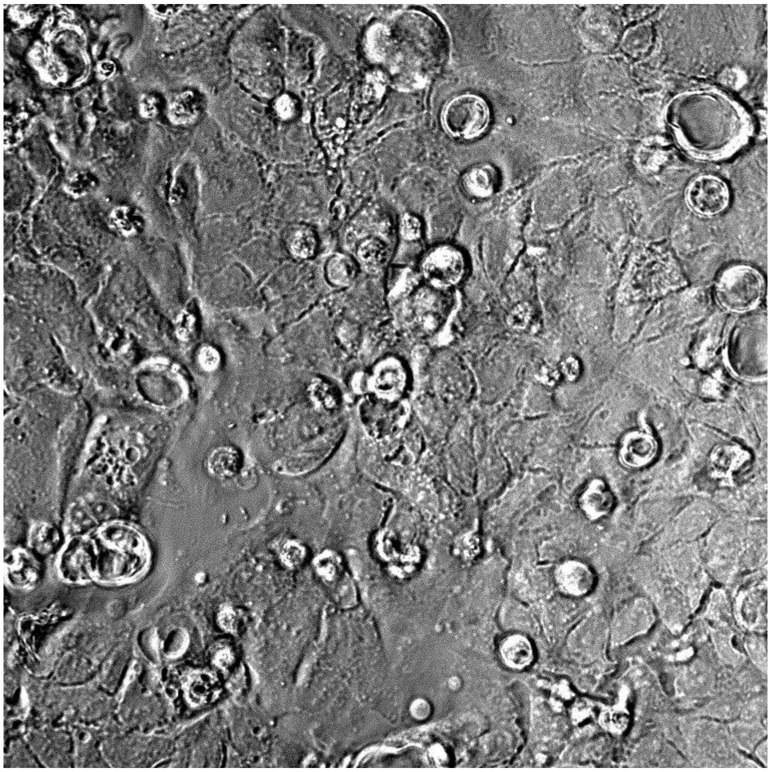

Caco-2, 1×10$^4$ cells/well, no oil, 24 h (bright field)

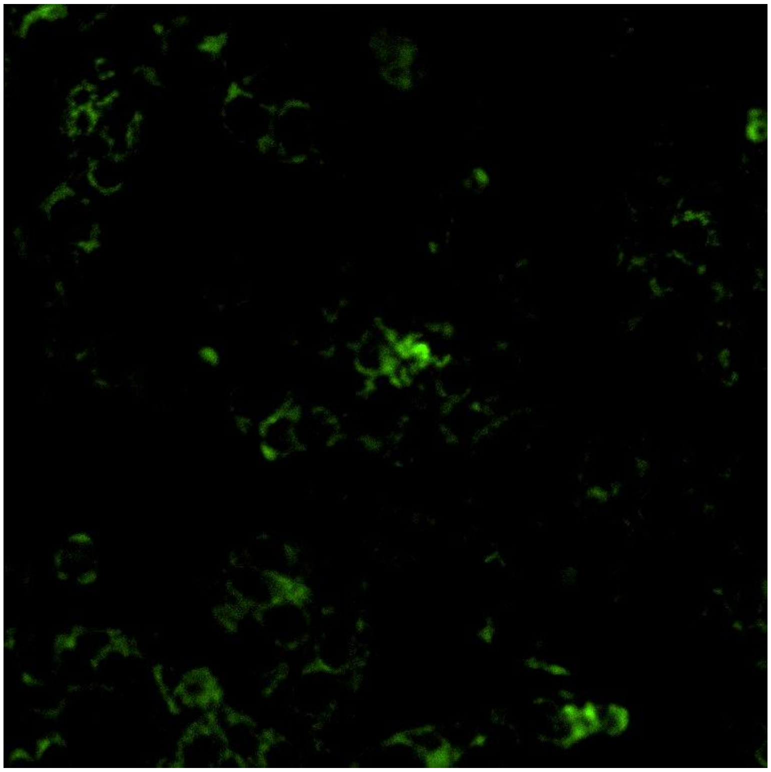

Caco-2, 1×10$^4$ cells/well, no oil, 24 h (hypoxia dye)

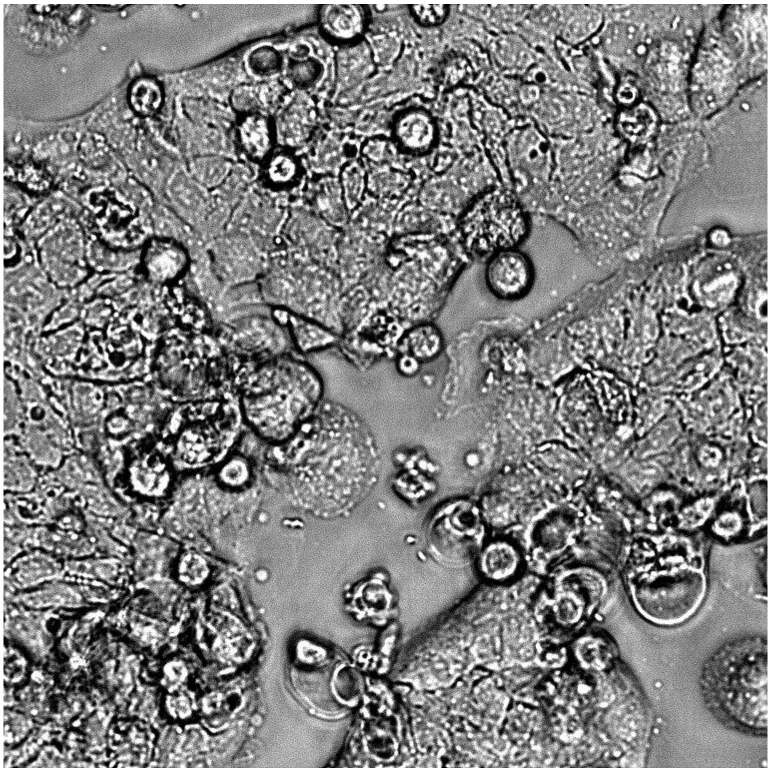

Caco-2, 1×10$^4$ cells/well, media-2 mm, silicone oil-5 mm-1000 cSt, 24 h (bright field)

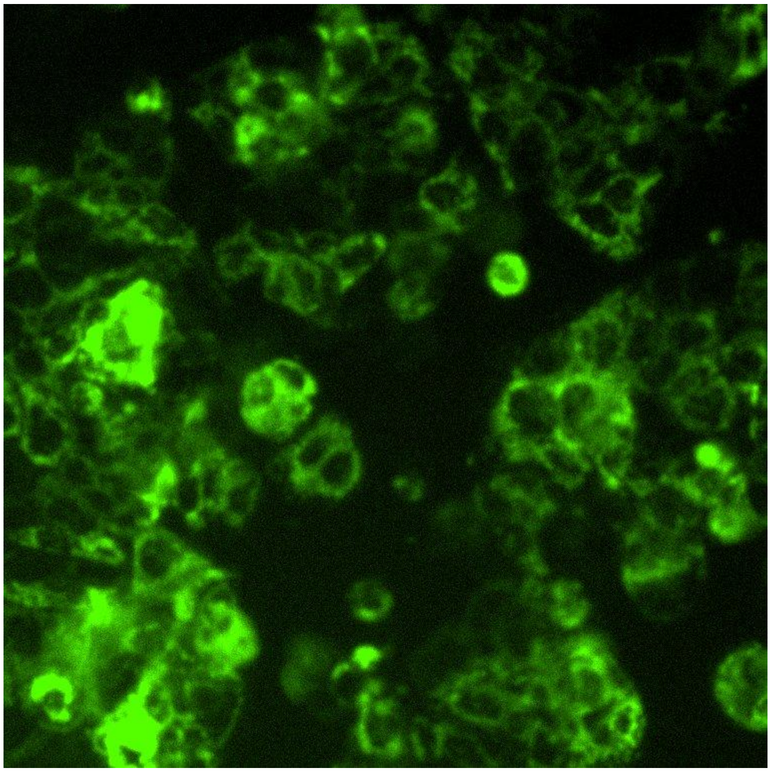

Caco-2, 1×10$^4$ cells/well, media-2 mm, silicone oil-5 mm-1000 cSt, 24 h (hypoxia dye)

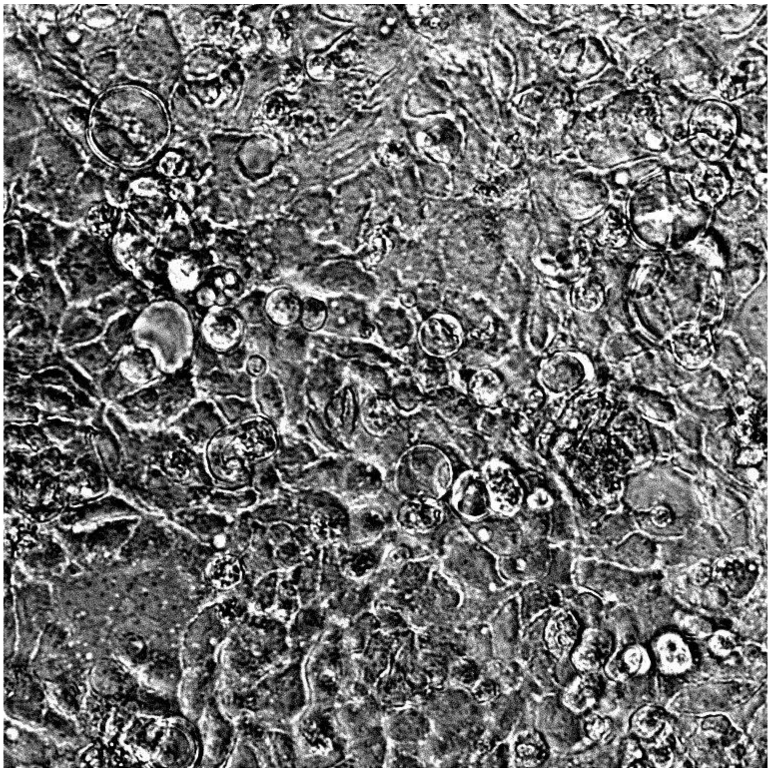

Caco-2, 3×10$^{4}$ cells/well, no oil, 24 h (bright field)

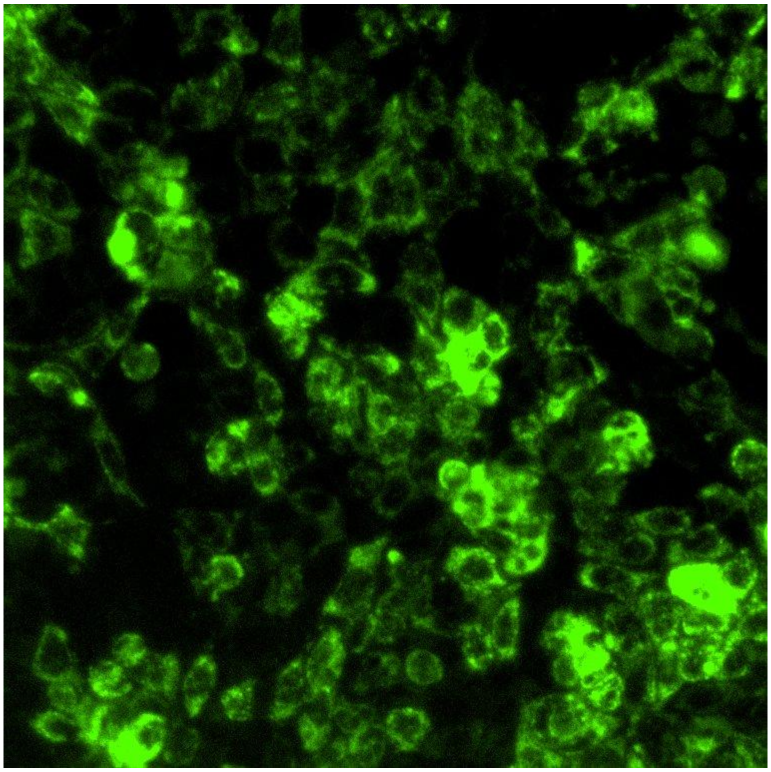

Caco-2, 3×10$^{4}$ cells/well, no oil, 24 h (hypoxia dye)

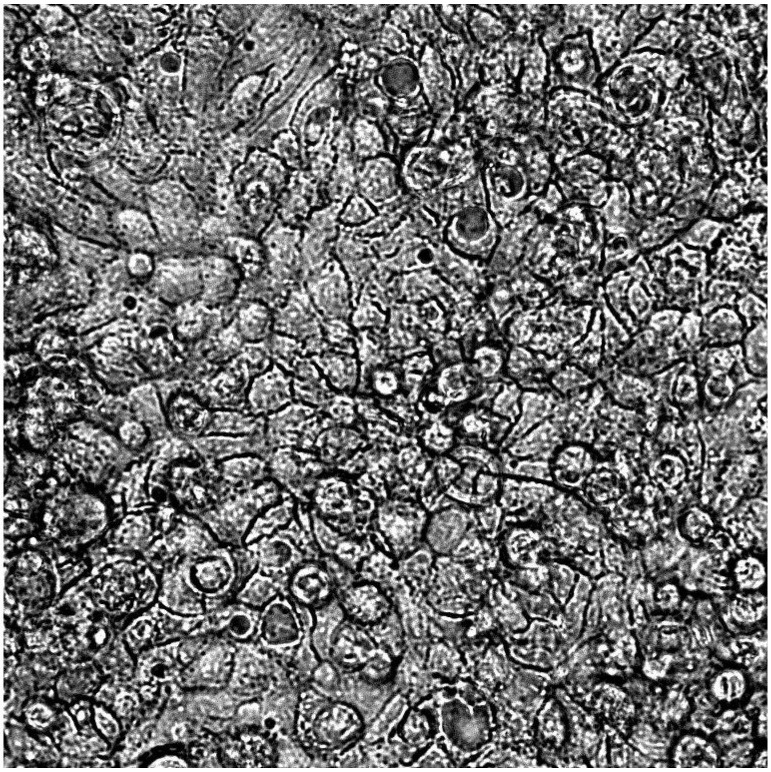

Caco-2, 3×$10^4$ cells/well, media-2 mm, silicone oil-5 mm-5 cSt, 24 h (bright field)

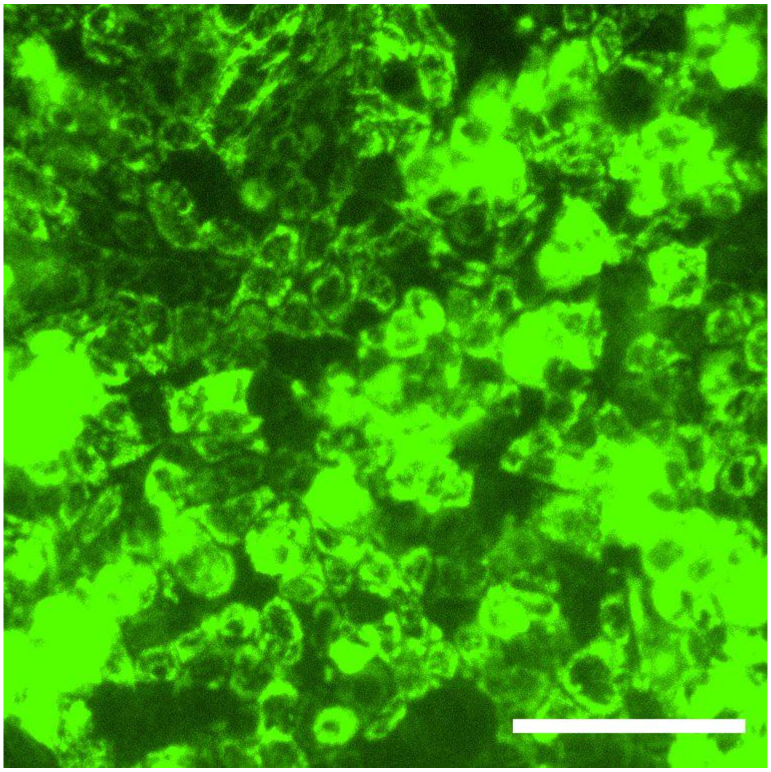

Caco-2, 3×$10^4$ cells/well, media-2 mm, silicone oil-5 mm-5 cSt, 24 h (hypoxia dye)

**Figure S7.** Large images showing the typical cell morphologies in Figure 4a. Scale bar, 100 µm.

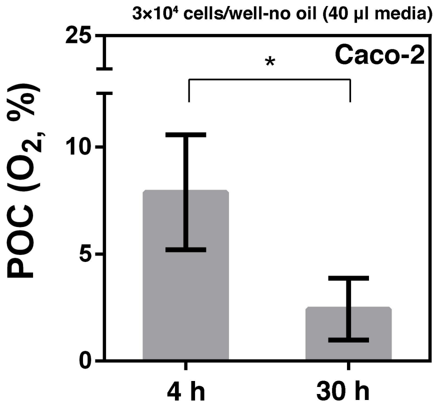


**Figure S8.** POC of Caco-2 from the condition of large media volume (40 µl/well on a 384-well plate for 4 mm in media depth) without oil overlay. Error bars, mean ± s.d. **P* ≤ 0.05.

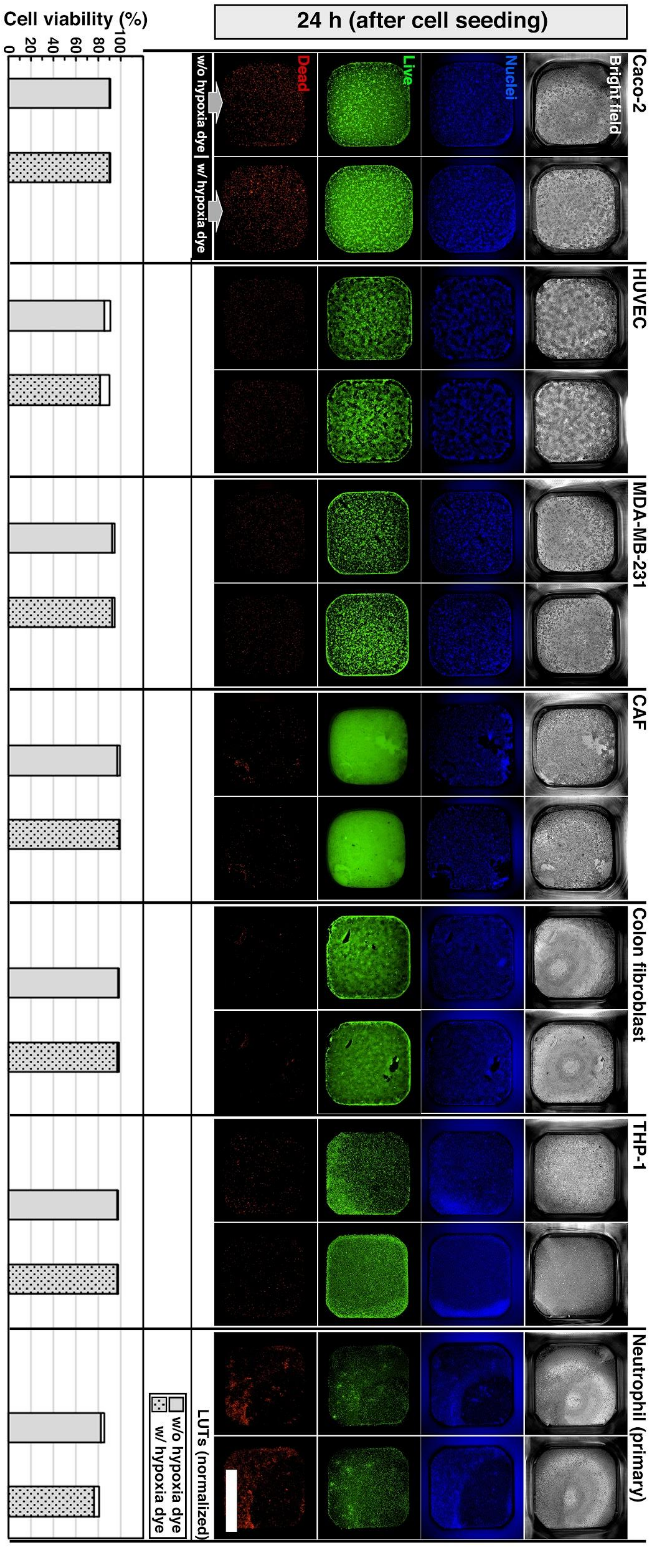


**Figure S9.** Cell viability of different cell types cultured under oil (24 h after cell seeding with and without hypoxia dye). Each cell type was seeded at $3{\times}10^4$ cells/well on a 384-well plate with 20 µl/well of media (for 2 mm in media depth), overlaid with 50 µl of silicone oil (5 cSt) (for 5 mm in oil depth), and cultured up to 24 h for a parallel comparison. The fluorescent images were all processed with normalized LUTs for visualization. Scale bar, 2 mm.

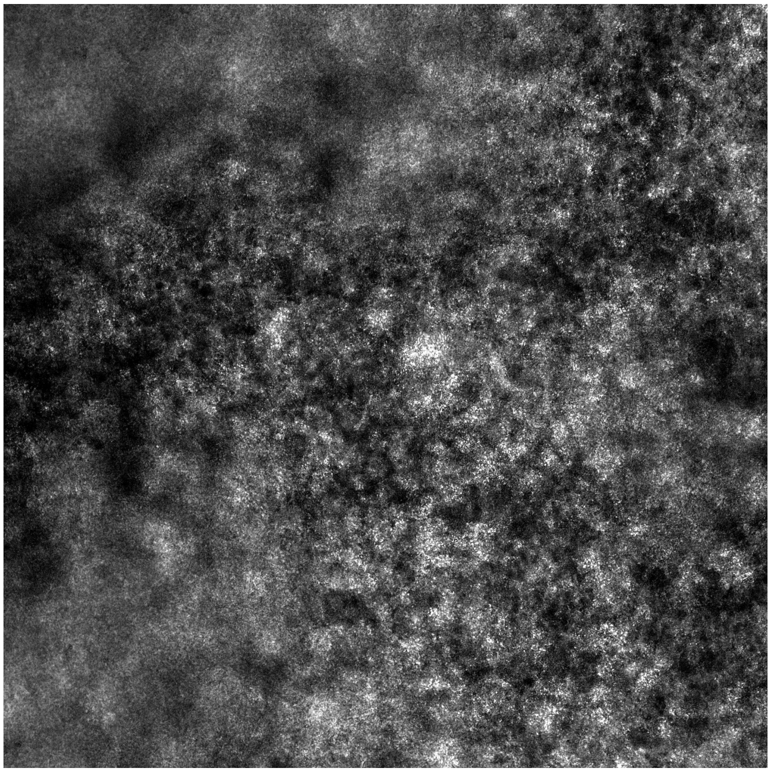

*C. albicans*, 3×$10^4$ cells/well, media-2 mm, silicone oil-5 mm-5 cSt, 24 h (bright field)

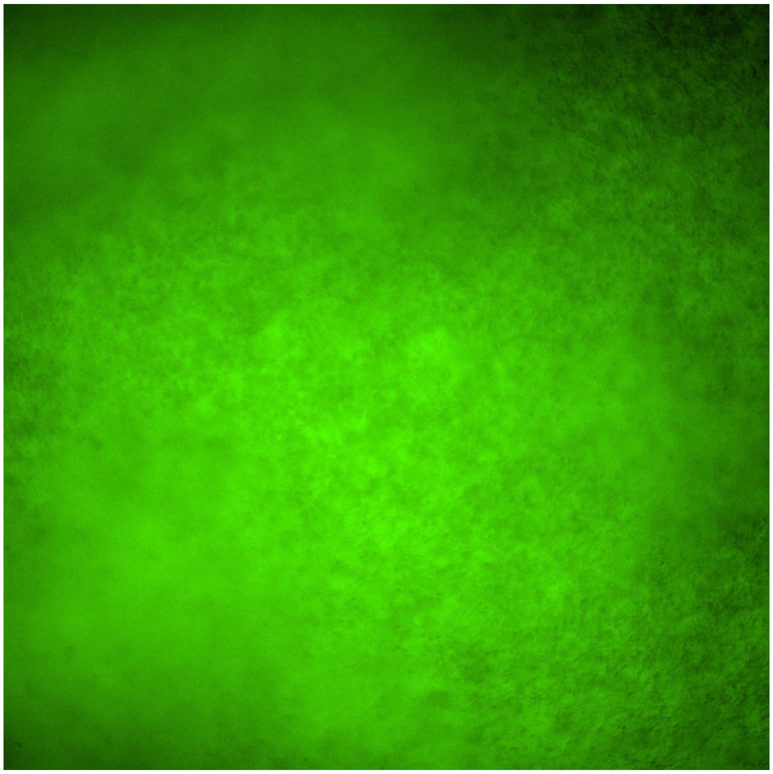

*C. albicans*, 3×$10^4$ cells/well, media-2 mm, silicone oil-5 mm-5 cSt, 24 h (hypoxia dye)

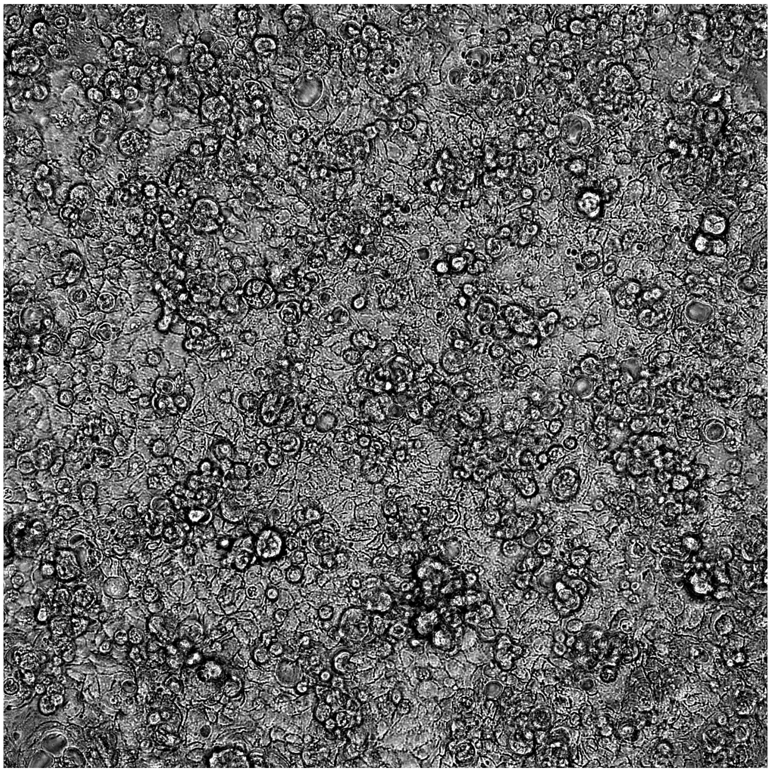

Caco-2, 3×10$^4$ cells/well, media-2 mm, silicone oil-5 mm-5 cSt, 24 h (bright field)

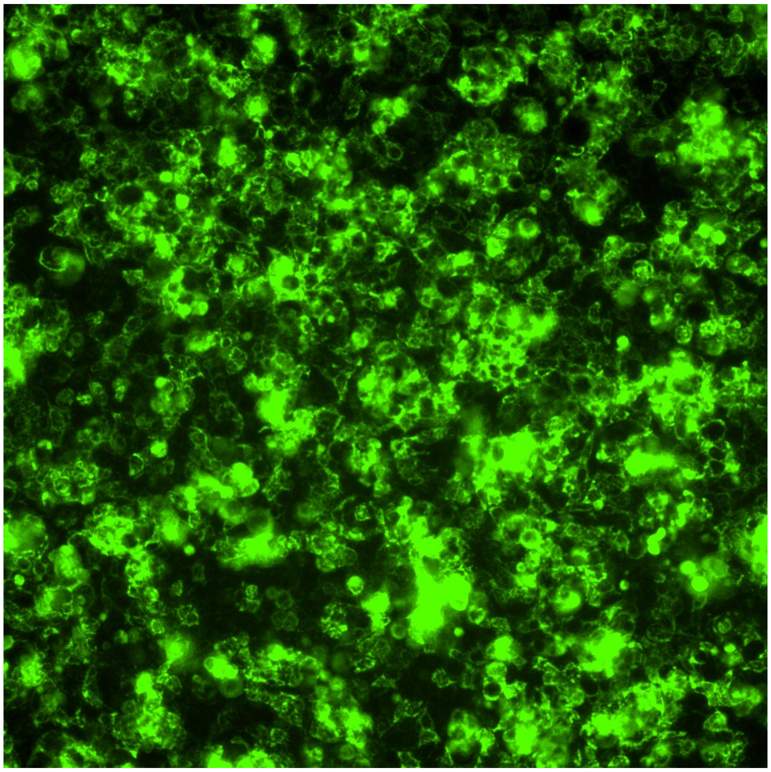

Caco-2, 3×10$^4$ cells/well, media-2 mm, silicone oil-5 mm-5 cSt, 24 h (hypoxia dye)

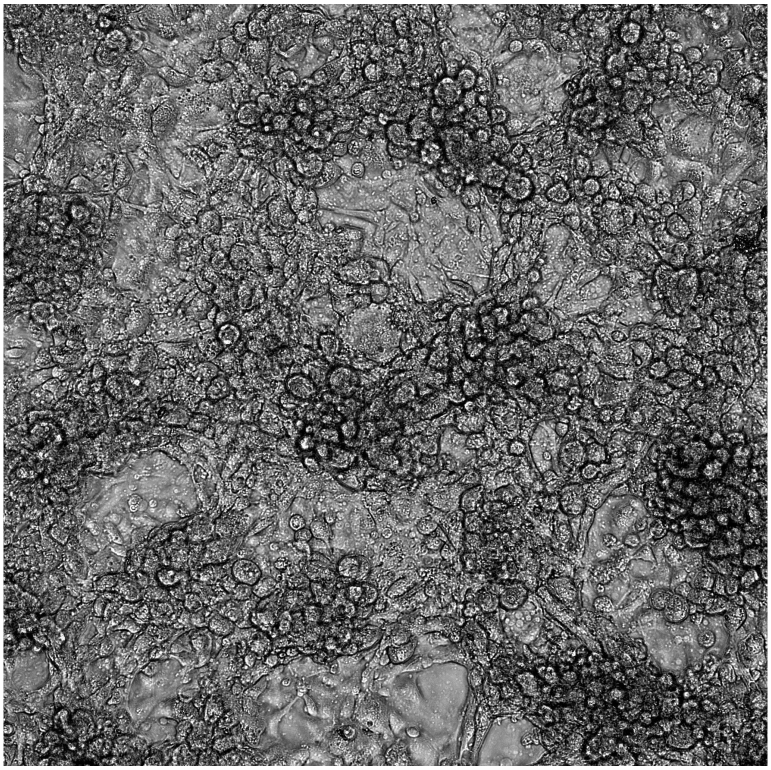
HUVEC, 3×$10^4$ cells/well, media-2 mm, silicone oil-5 mm-5 cSt, 24 h (bright field)

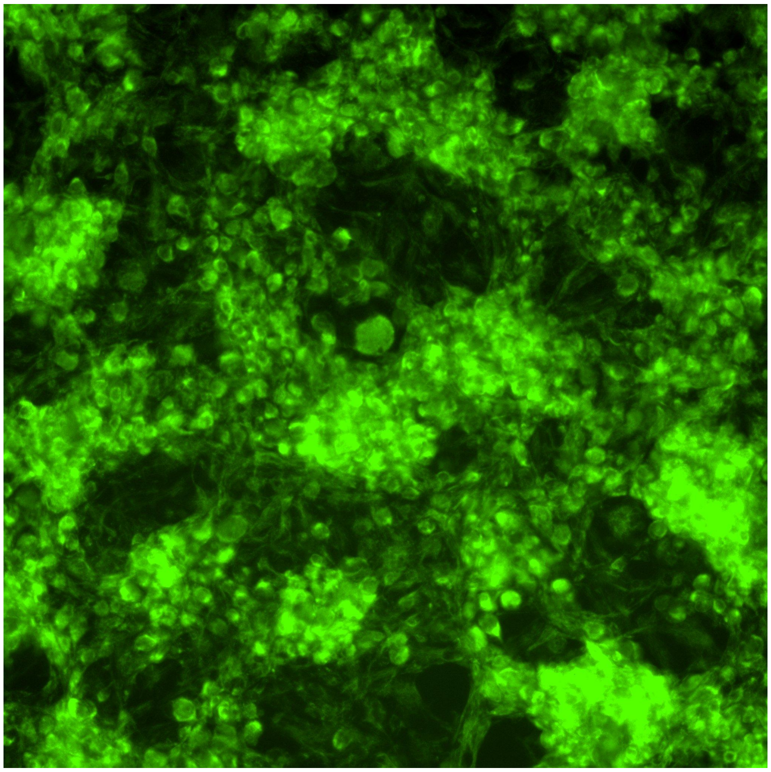
HUVEC, 3×$10^4$ cells/well, media-2 mm, silicone oil-5 mm-5 cSt, 24 h (hypoxia dye)

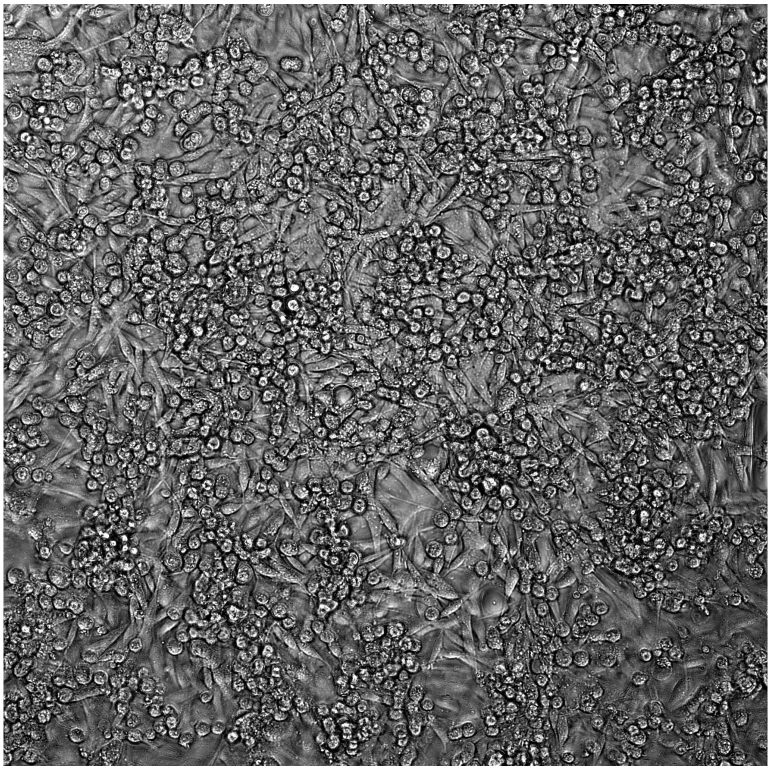

MDA-MB-231, $3\times10^4$ cells/well, media-2 mm, silicone oil-5 mm-5 cSt, 24 h (bright field)

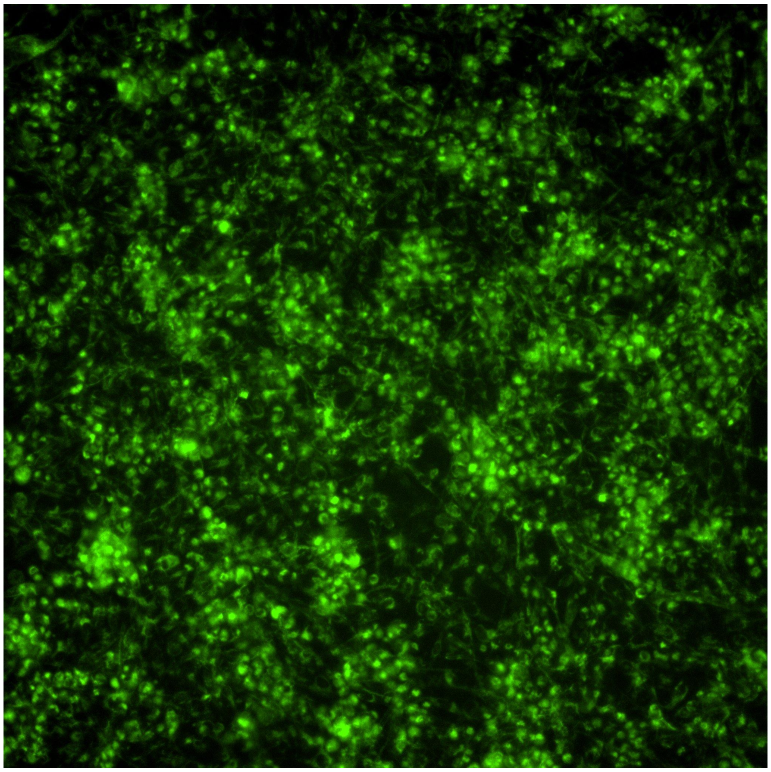

MDA-MB-231, $3\times10^4$ cells/well, media-2 mm, silicone oil-5 mm-5 cSt, 24 h (hypoxia dye)

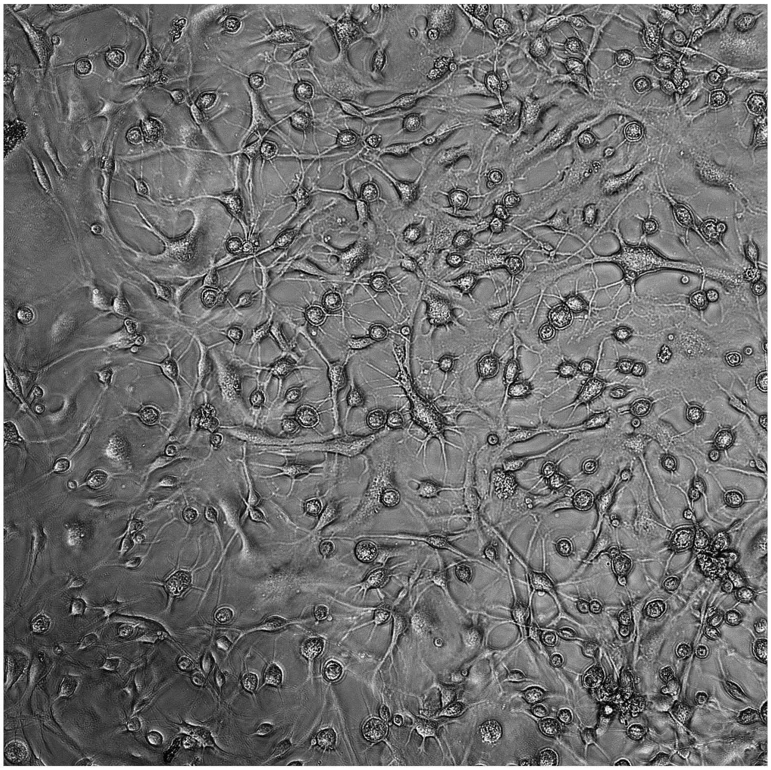
CAF, 3×$10^4$ cells/well, media-2 mm, silicone oil-5 mm-5 cSt, 24 h (bright field)

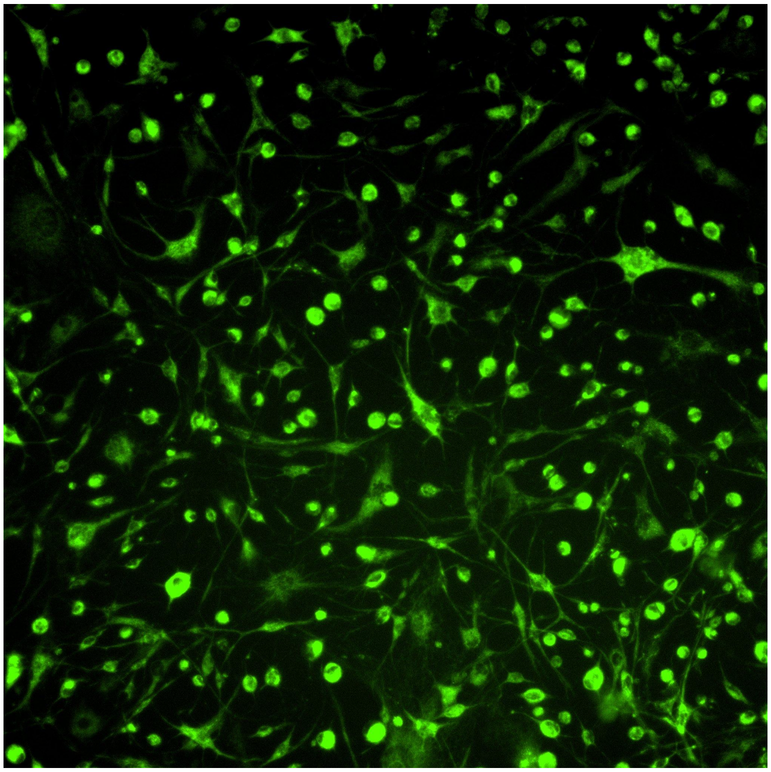
CAF, 3×$10^4$ cells/well, media-2 mm, silicone oil-5 mm-5 cSt, 24 h (hypoxia dye)

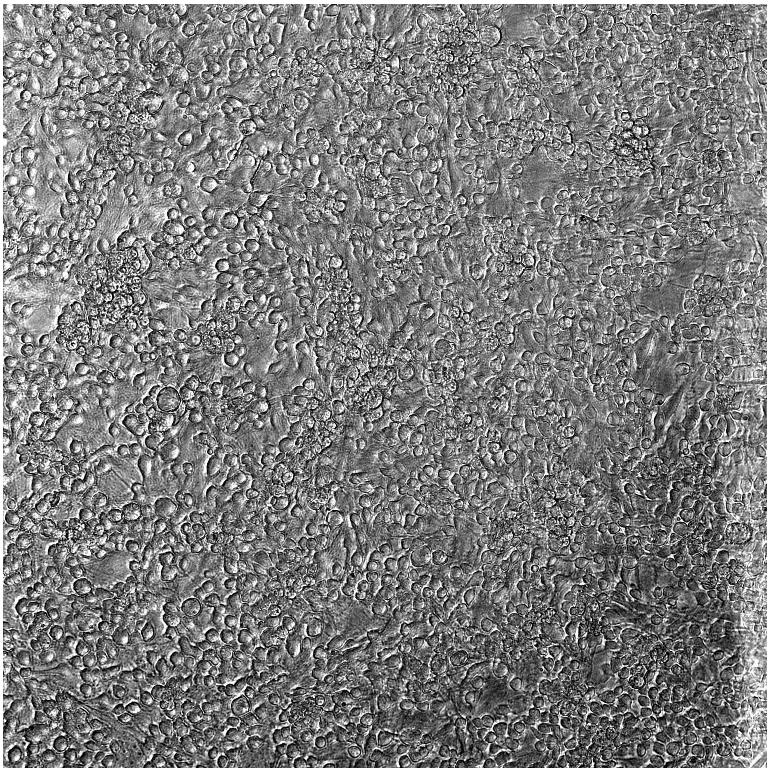

Colon fibroblast, 3×10$^4$ cells/well, media-2 mm, silicone oil-5 mm-5 cSt, 24 h (bright field)

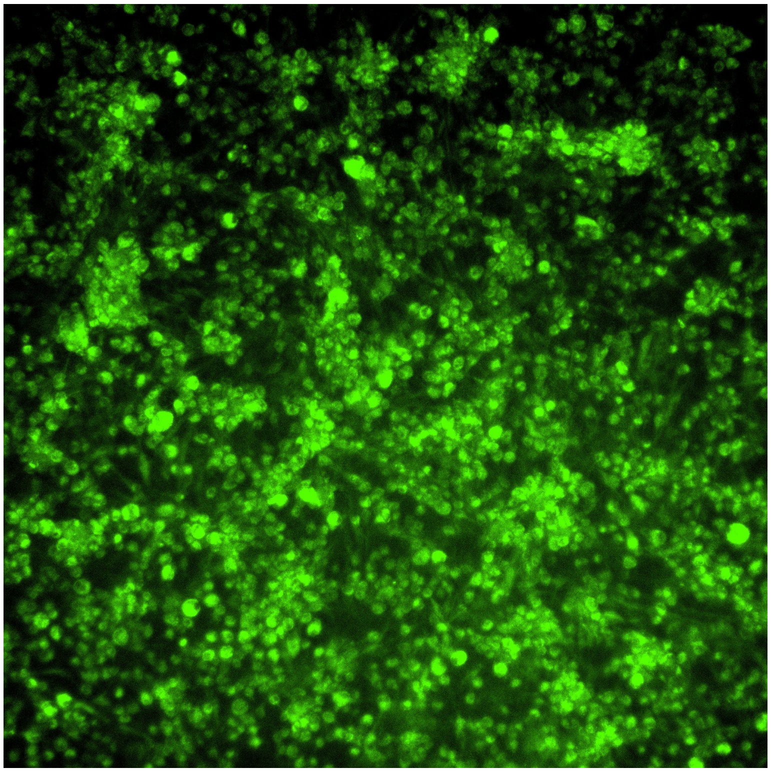

Colon fibroblast, 3×10$^4$ cells/well, media-2 mm, silicone oil-5 mm-5 cSt, 24 h (hypoxia dye)

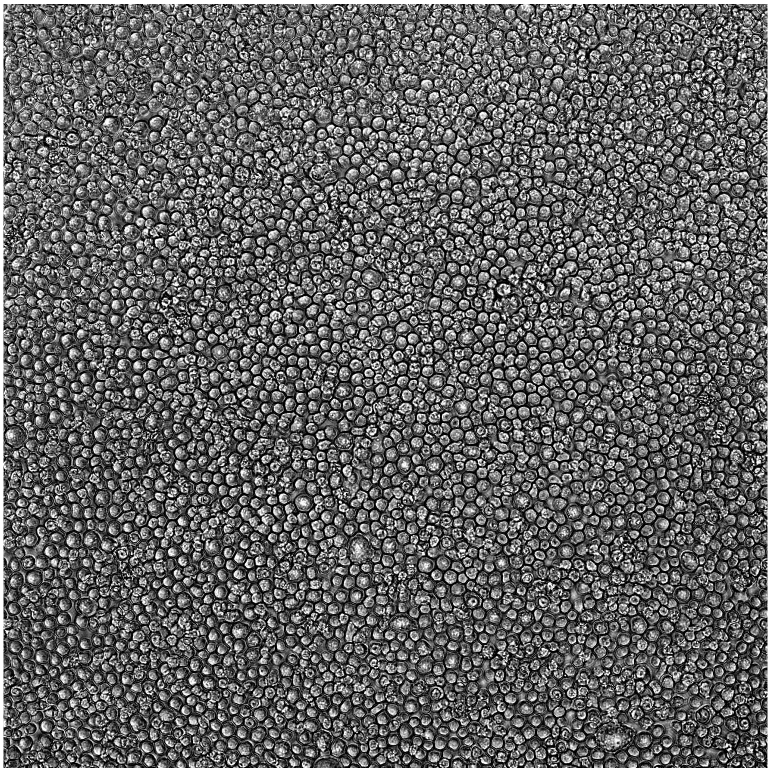

THP-1, 3×10$^4$ cells/well, media-2 mm, silicone oil-5 mm-5 cSt, 24 h (bright field)

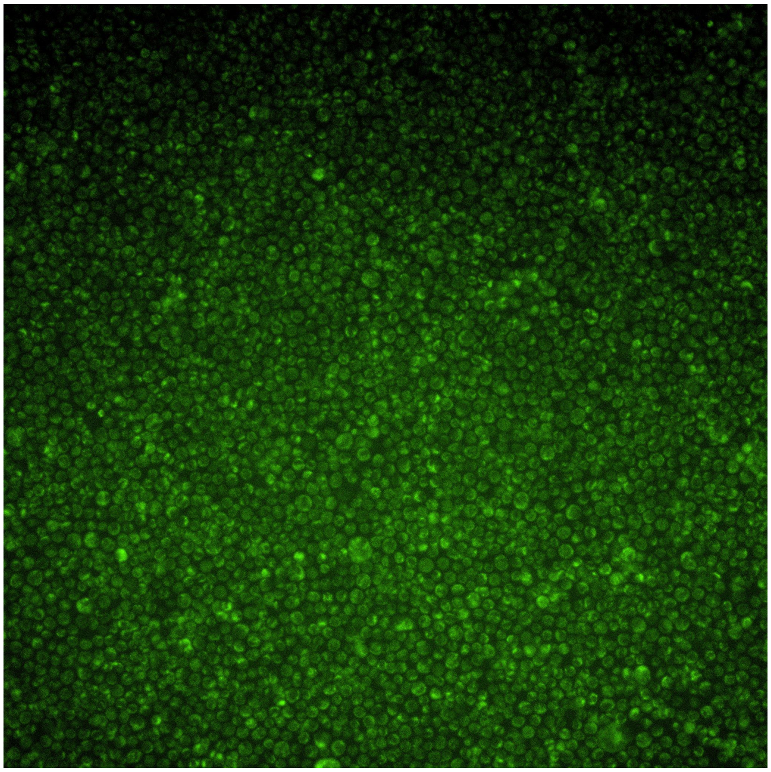

THP-1, 3×10$^4$ cells/well, media-2 mm, silicone oil-5 mm-5 cSt, 24 h (hypoxia dye)

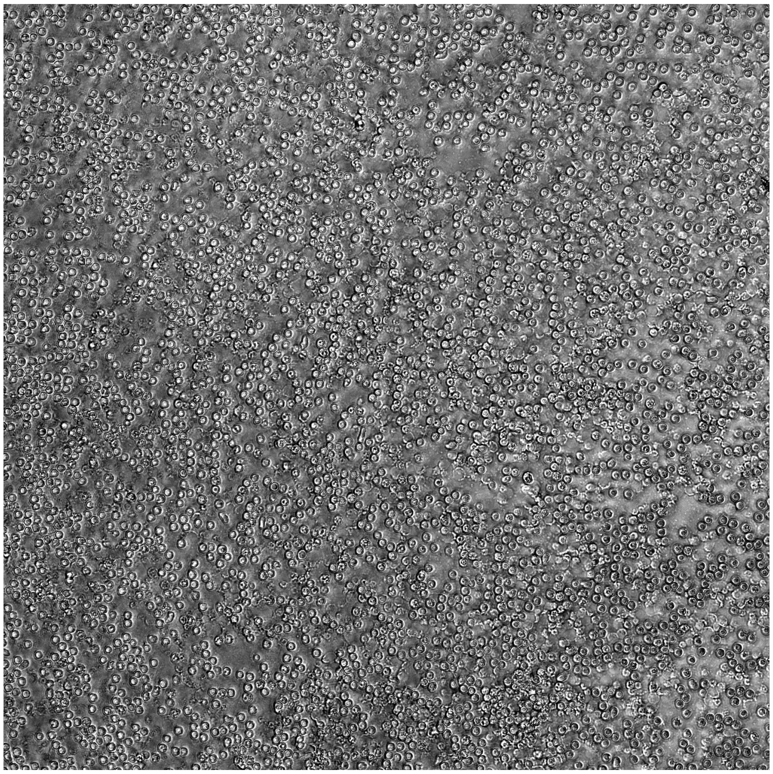

Neutrophil, 3×10$^4$ cells/well, media-2 mm, silicone oil-5 mm-5 cSt, 24 h (bright field)

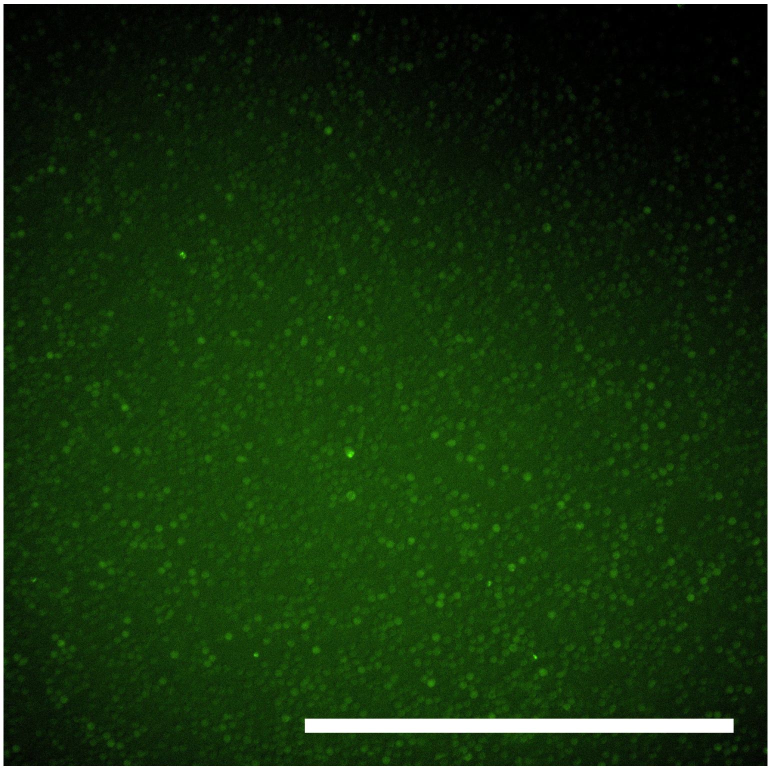

Neutrophil, 3×10$^4$ cells/well, media-2 mm, silicone oil-5 mm-5 cSt, 24 h (hypoxia dye)

**Figure S10.** Large images showing the typical cell morphologies in Figure 5b. Scale bar, 500 μm.

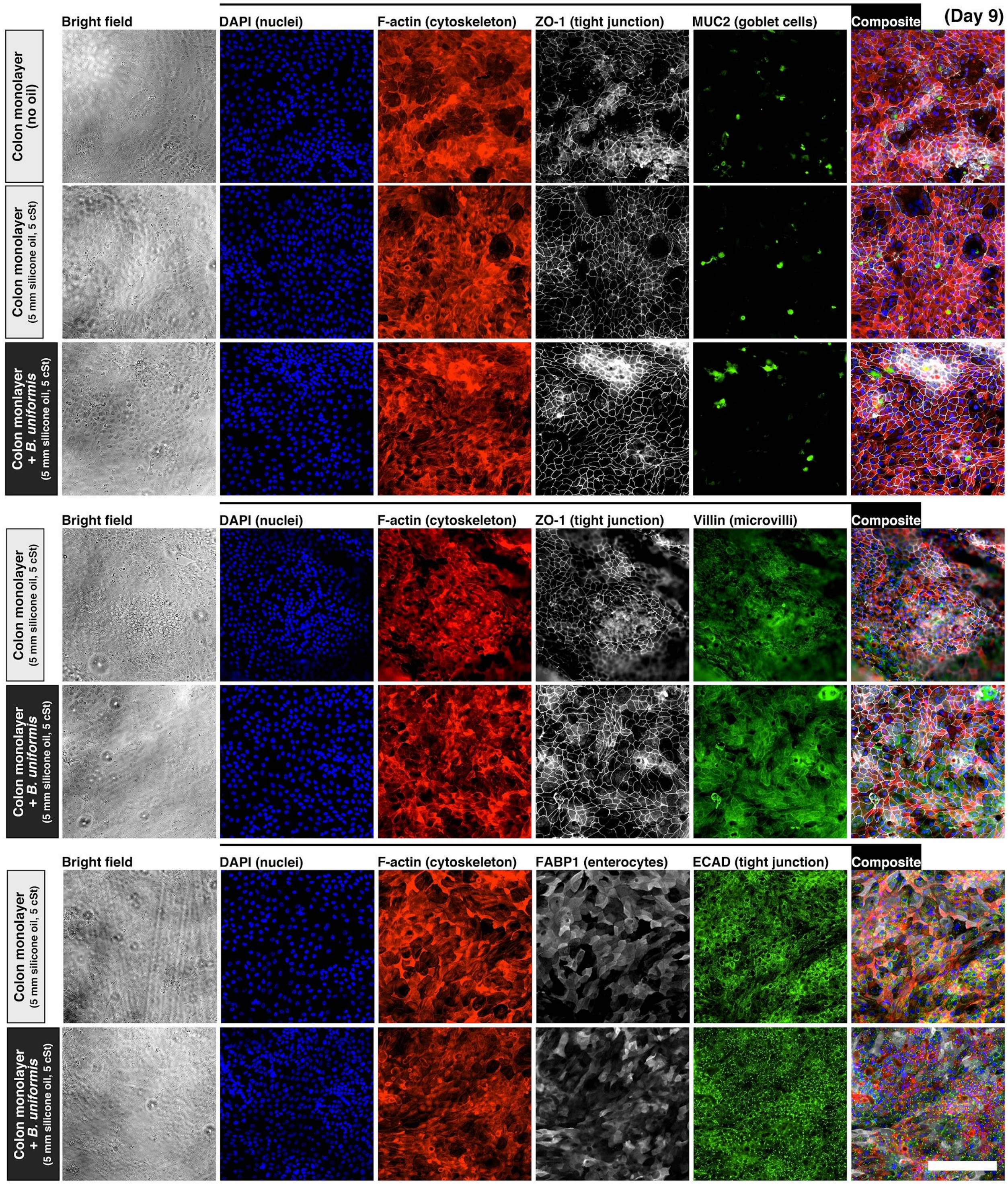


**Figure S11.** IFS images of primary colon epithelium from mono-culture (no-bacteria control) and co-culture with *B. uniformis* under oil on Day 9 (i.e. 24 h after inoculation of the bacteria). The fluorescent images were all processed with normalized LUTs for visualization. Scale bar, 200 μm.

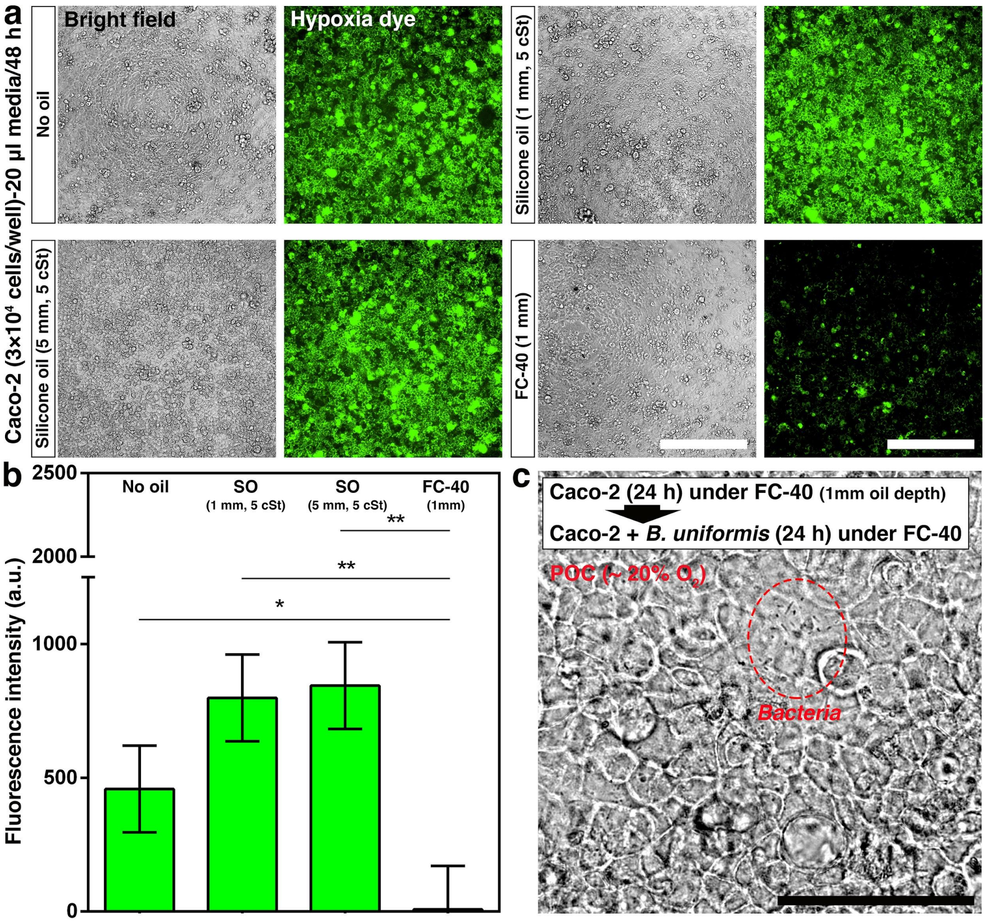


**Figure S12.** Comparison of hypoxia generation between silicone oil (5 cSt) and fluorinated oil (Fluorinert FC-40). a) Microscopic images (bright field, left; fluorescent, right) of Caco-2 monolayers [$3\times10^4$ cells/well, 20 µl/well of media (for 2 mm media depth), 10 or 50 µl/well silicone oil (5 cSt) (for 1 or 5 mm oil depth, respectively) overlay, 10 µl/well fluorinated oil (FC-40) (for 1 mm in oil depth)] cultured on a 384-well plate for 48 h. The fluorescent images of hypoxia dye were processed with parallel LUTs. Scale bars, 500 µm. b) IOC (fluorescence intensity of hypoxia dye) of each condition. Error bars, mean ± s.d. *$P \leq 0.05$, and **$P \leq 0.01$. c) Co-culture of Caco-2 monolayer from (a) with *B. uniformis* [inoculum density, $OD_{600}$ = 0.1, 1:20 v/v ratio (1 µl bacteria:20 µl media)] under fluorinated oil (FC-40, 1 mm oil depth). POC was measured for about 20% $O_2$ with the fluorinated oil overlay. The bacteria (the red dashed line circle) showed little growth after 24 h co-culture under FC-40. Scale bar, 200 µm.

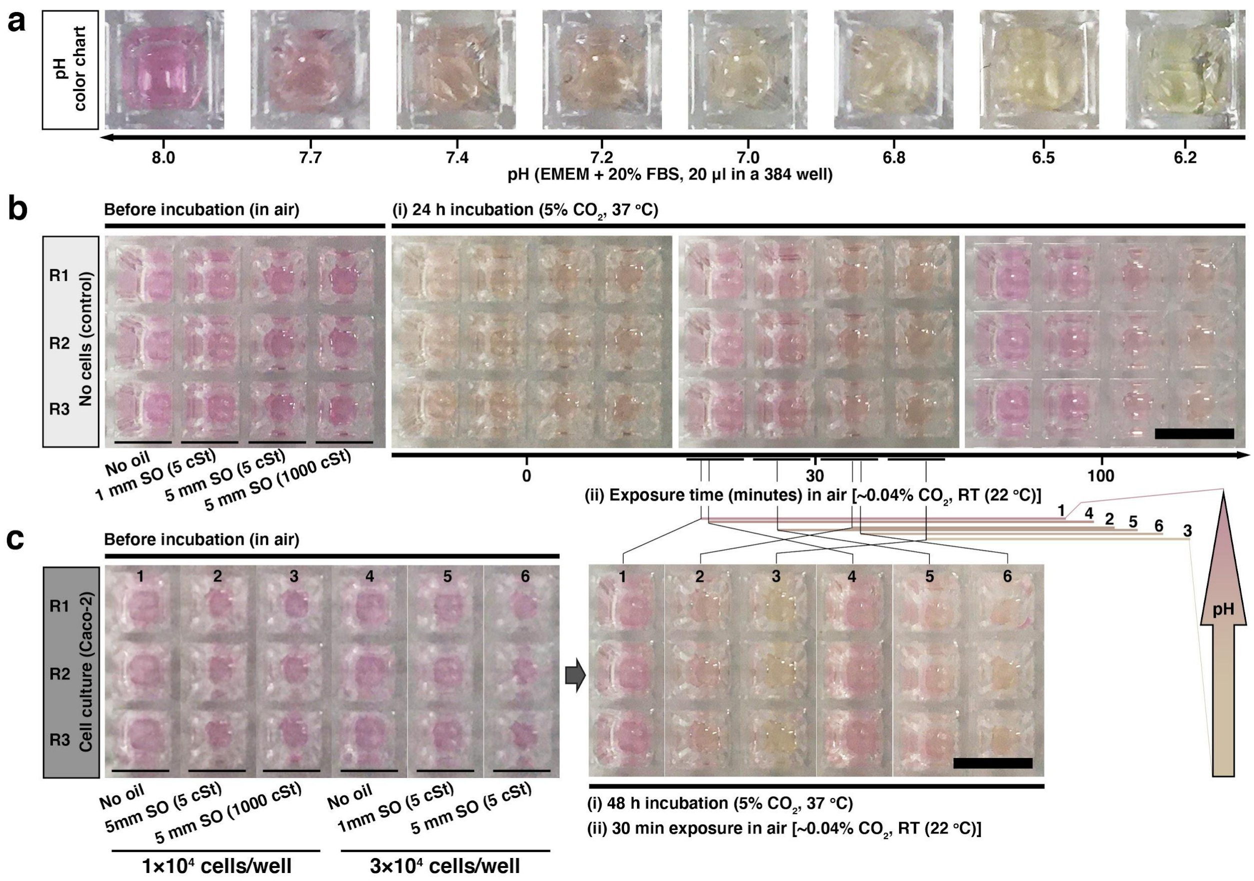


**Figure S13.** Colorimetric analysis of pH of the culture media before, after incubation and exposure in air. a) A pH color chart of phenol red (30 µM in EMEM + 20% FBS, 20 µl/well on a 384-well plate). b) The control of a no-cell plate with different oil [silicone oil (SO)] overlays. c) The Caco-2 plate [1×10$^4$ or 3×10$^4$ cells/well, 20 µl/well of media (for 2 mm in media depth)] with different oil overlays. $CO_2$ dissolved in the culture media diffused out through the oil overlay over time, which led to the different recovery rates of pH to basic across the tested conditions. The oil overlay stabilized the pH in culture media during device transfer or operation in an atmospheric ambient environment. Scale bars, 4 mm.

[1] K. L. VanDussen, N. M. Sonnek, T. S. Stappenbeck, *Stem Cell Res.* **2019**, *37*, 101430.
[2] M. M. Mahe, N. Sundaram, C. L. Watson, N. F. Shroyer, M. A. Helmrath, *J. Vis. Exp.* **2015**, DOI 10.3791/52483.
